# Discovery and prioritization of variants and genes for kidney function in >1.2 million individuals

**DOI:** 10.1101/2020.09.04.283713

**Authors:** Kira J Stanzick, Yong Li, Mathias Gorski, Matthias Wuttke, Cristian Pattaro, Anna Köttgen, Klaus J Stark, Iris M Heid, Thomas W Winkler

## Abstract

Chronic kidney disease (CKD) has a complex genetic underpinning. Genome-wide association studies (GWAS) of CKD-defining glomerular filtration rate (GFR) have identified hundreds of loci, but prioritization of variants and genes is challenging. To expand and refine GWAS discovery, we meta-analyzed GWAS data for creatinine-based estimated GFR (eGFRcrea) from the Chronic Kidney Disease Genetics Consortium (CKDGen, n=765,348, trans-ethnic) and UK Biobank (UKB, n=436,581, Europeans). The results (i) extend the number of eGFRcrea loci (424 loci; 201 novel; 8.9% eGFRcrea variance explained by 634 independent signals); (ii) improve fine-mapping resolution (138 99% credible sets with ≤5 variants, 44 single-variant sets); (iii) ascertain likely kidney function relevance for 343 loci (consistent association with alternative biomarkers); and (iv) highlight 34 genes with strong evidence by a systematic Gene PrioritiSation (GPS). We provide a sortable, searchable and customizable GPS tool to navigate through the *in silico* functional evidence and select relevant targets for functional investigations.

## INTRODUCTION

Chronic kidney disease (CKD) is a leading cause of morbidity and mortality worldwide, and a major public health problem with prevalence of >10% in the adult population in developed countries ^1,2^. Although many underlying causes of CKD such as diabetes, vascular disease, or glomerulonephritis are known, CKD etiology remains in most cases unclear. Moreover, knowledge about the underlying molecular mechanisms causing progressive loss of renal function is so far insufficient, resulting in a lack of therapeutic targets for drug development ^3^.

A hallmark of CKD is decreased glomerular filtration rate, which can be estimated from the serum creatinine level ^4^. Estimated creatinine-based GFR (eGFRcrea) has a strong heritable component ^5^. Twin studies estimated a broad-sense heritability for eGFRcrea of 54% ^5^. Recently, a GWAS meta-analysis of eGFRcrea conducted by the CKD Genetics (CKDGen) Consortium identified 264 associated genetic loci ^6,7^. The index SNPs at the identified loci explained nearly 20% of eGFRcrea’s genetic heritability ^7^. While eGFRcrea is a useful marker of kidney function in clinical practice, the underlying serum creatinine is a metabolite from muscle metabolism ^8,9^ and thus may not only reflect kidney function. Alternative kidney function biomarkers such as GFR estimated by serum cystatin C (eGFRcys) and blood urea nitrogen (BUN) have had, so far, a more limited role in large population-based studies and GWAS ^7,10^. On the genetic side, a major challenge is that genetic association loci highlight the region with the best statistical evidence for association but they provide only limited insights into the causal genes, variants or biological mechanisms. Numerous approaches for bioinformatic functional characterization of identified loci yield an abundance of potentially relevant information, but the more the identified loci, the larger the jungle of evidence ^11–13^.

We thus conducted a meta-analysis of GWAS to increase our knowledge of genetic loci associated with kidney function using data from >1.2 million individuals from the CKDGen Consortium ^7^ and UK Biobank (UKB) ^14^, integrated genetic data for eGFRcys and BUN in >400,000 individuals, and submitted the identified loci to a systematic functional bioinformatic follow-up analysis. A major aim of our effort was to provide a customizable and searchable overview for the abundance of results to enable the prioritization of genes and variants, which should help select relevant targets for functional follow-up. The workflow is illustrated in **Supplementary Figure 1**.

## RESULTS

### GWAS meta-analysis identified 201 novel non-overlapping loci for eGFRcrea

To identify genetic variants associated with eGFRcrea, we conducted a linear mixed model-based GWAS ^15^ of eGFRcrea in UKB (European ancestry, n = 436,581, mean +/− SD age = 56.8 +/− 8.0 years, mean +/− SD eGFRcrea = 90.5 +/− 13.3 ml/min/1.73m^2^, imputed to Haplotype Reference Consortium ^16^ and UK10K panels ^17^) and meta-analyzed results with the CKDGen Consortium data (trans-ethnic, n = 765,348, imputed to Haplotype Reference Consortium ^16^ or 1000 Genomes ^18^) ^7^, for a total sample size of 1,201,909 individuals (**Supplementary Figure 1, Methods**). From the 13,633,840 variants with a minor allele frequency (MAF) of >=0.1%, we selected genome-wide significant variants (GWS, P < 5 x 10^−8^) and derived non-overlapping loci using a stepwise approach (locus region defined by the first and last GWS variant of a locus +/−250kb, **Methods**).

We identified 424 non-overlapping loci: 201 were novel and 223 were known (**Figure 1**, **Supplementary Table 1, Supplementary Figure 2**). We considered known a locus with at least one GWS variant included within one of the 264 loci previously identified by Wuttke *et al* ^7^ (**Methods**). Only three of the 264 loci from Wuttke *et al.* ^7^ barely missed genome-wide significance in our meta-analysis, which can be attributed to chance (P < 7.5 x 10^−7^, **Supplementary Table 2**). We observed more low-frequency variants among novel compared to known loci (7.0% versus 2.2% variants had MAF <5% among novelversus known, **Figure 1B**).

**Figure 1.**
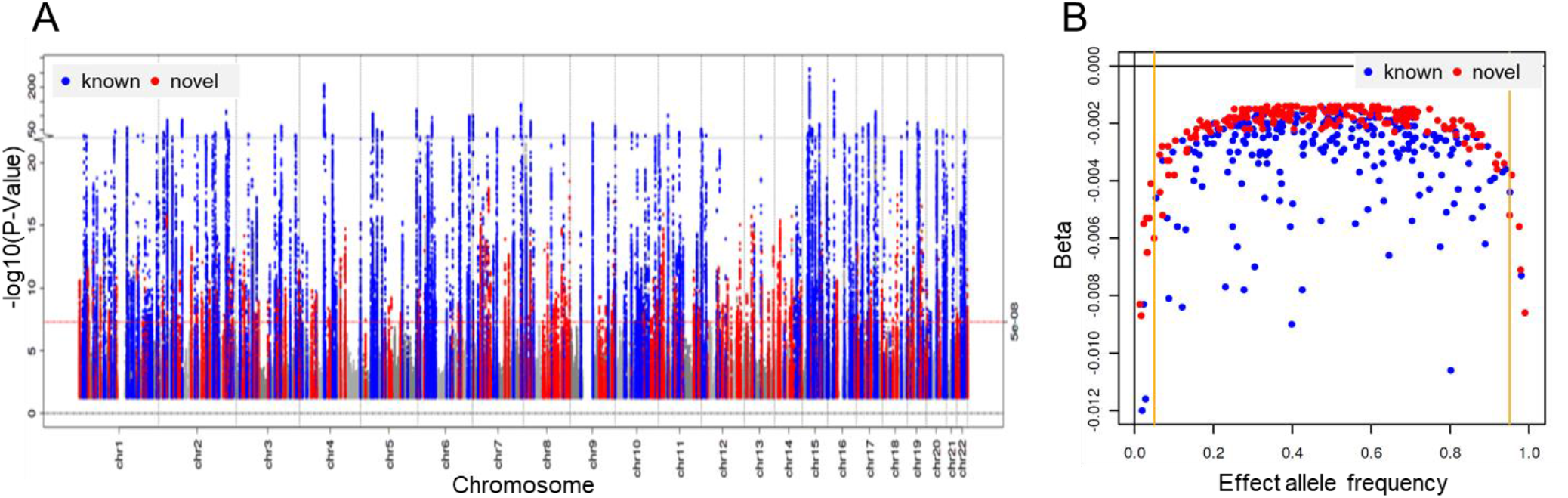
Primary meta-analysis results for eGFRcrea. Shown are results from our primary meta-analysis for eGFRcrea (n = 1,201,929). A: Manhattan-Plot showing –log10 association P value for the genetic effect on eGFRcrea by chromosomal base position (GRCh37, 223 known loci are marked in blue, 201 novel loci in red). The red dashed line marks genome-wide significance (5 x 10^−8^). B: Scatter plot comparing eGFRcrea effect sizes versus allele frequencies for the 424 identified locus lead variants. Effect sizes and allele frequencies were aligned to the eGFRcrea decreasing alleles. Coloring is analogous to A. Orange lines mark allele frequencies of 5% and 95%.

To evaluate the impact of the 197,888 individuals of non-European ancestry (CKDGen) on our primary meta-analysis results, we conducted a sensitivity meta-analysis limited to individuals of European ancestry (CKDGen, n = 567,460; UKB as before; total n = 1,004,040, **Methods**). The genetic effect sizes for the 424 lead variants showed highly consistent results between the primary and the European-only analysis (**Supplementary Table 3, Supplementary Figure 3**), suggesting lack of between-ancestry heterogeneity for the identified variants.

Taken together, the meta-analysis of CKDGen and UKB identified 424 independent, non-overlapping loci for eGFRcrea, including 201 novel and 223 known loci. In the following, these loci are evaluated with regard to alternative kidney function biomarker, secondary signals, fine-mapping and gene prioritization.

### Association of identified variants with alternative kidney function biomarkers

A genetic association with eGFRcrea can be related to kidney function or to creatinine metabolism. We thus evaluated the 424 lead variants for their association with two other kidney-related traits. We conducted GWAS of eGFRcys and BUN in UKB and meta-analysed results with data from the CKDGen Consortium ^7,10^ (combined n = 460,826 for eGFRcys, n = 852,678 for BUN, **Methods**). We defined an eGFRcrea association as validated by eGFRcys or BUN when we observed a directionally consistent nominally significant association with eGFRcys and/or BUN (i.e. same effect direction for eGFRcys or opposite effect direction for BUN). Of the 424 lead variants, 132 were consistent only for eGFRcys, 23 only for BUN and 188 for both (**Figure 2, Supplementary Table 4**). Despite a doubling of sample size for BUN vs. Wuttke et al ^7^, we found a similar proportion of BUN-validated eGFRcrea loci (211 of 424 loci, 50%; compared to 146 of 264 Wuttke et al. loci, 55%). Even with much lower sample size, we found 320 eGFRcys validated loci (of 424 loci, 75%).

In summary, ~81% (343 loci) of the identified eGFRcrea loci were validated by at least one alternative biomarker and thus classified as likely relevant for kidney function.

**Figure 2.**
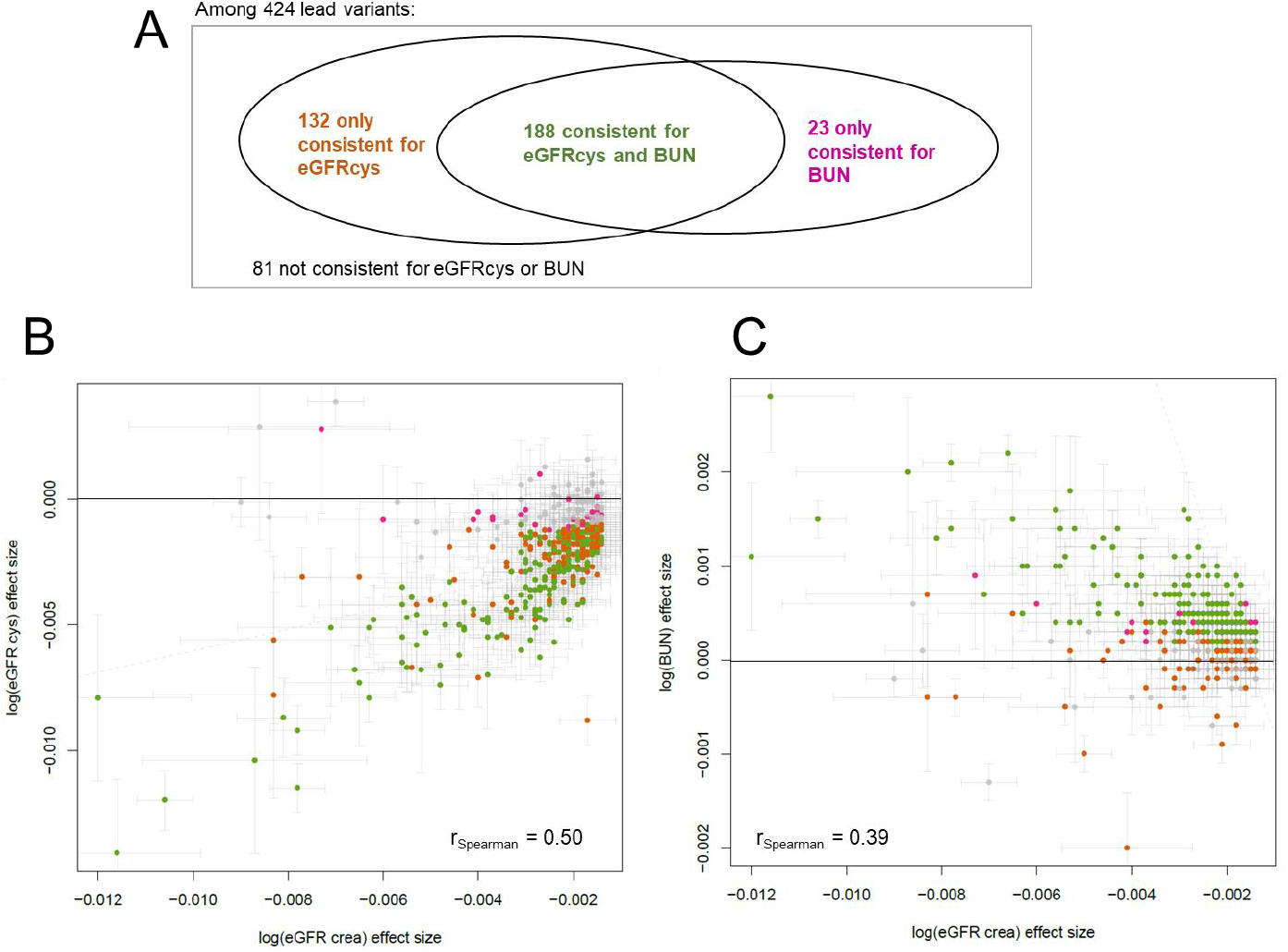
Relevance for kidney function based on alternative biomarker analyses. Shown are results from our evaluation of kidney function relevance for the 424 locus lead variants identified by our primary meta-analysis for eGFRcrea (n = 1,201,929). We classified the 424 variants by their nominal significantsignificant (P<0.05) and consistent effect direction for BUN (n = 679,531, i.e. opposite effect to eGFRcrea) or eGFRcys (n = 460,826, i.e. same effect direction as eGFRcrea). A: Venn diagram showing the distribution of 424 lead variants by alternative biomarker relevance. B: Scatterplot comparing effect sizes for eGFRcrea and eGFRcys (green: eGFRcys and BUN validated, brown: only eGFRcys validated, magenta: only BUN validated, grey: not validated). C: Scatterplot comparing effect sizes for eGFRcrea and BUN (coloring analogous to B).

### Secondary signals and fine-mapping in European ancestry

To identify multiple independent variant associations, i.e. signals, at the 424 non-overlapping loci, we conducted conditional analyses using GCTA ^19^ (**Methods**). Due to the lack of an appropriate trans-ethnic linkage disequilibrium (LD) reference panel, we here focused on the European-only results and used a random subset of 20,000 unrelated individuals of European-ancestry from UKB as LD reference panel (**Methods**). We identified 634 independent signals (P-value conditioned on other signal lead variants < 5 x 10^−8^) across the 424 loci (**Figure 3A**, **Supplementary Table 5**). At least two independent signals were observed at 21 novel (**Supplementary Figure 4**) and 101 known loci. In the known *UMOD/PDILT* locus, we observed four independent signals, two novel and two previously described ^7^ (**Supplementary Figure 5, Supplementary Table 5**).

**Figure 3.**
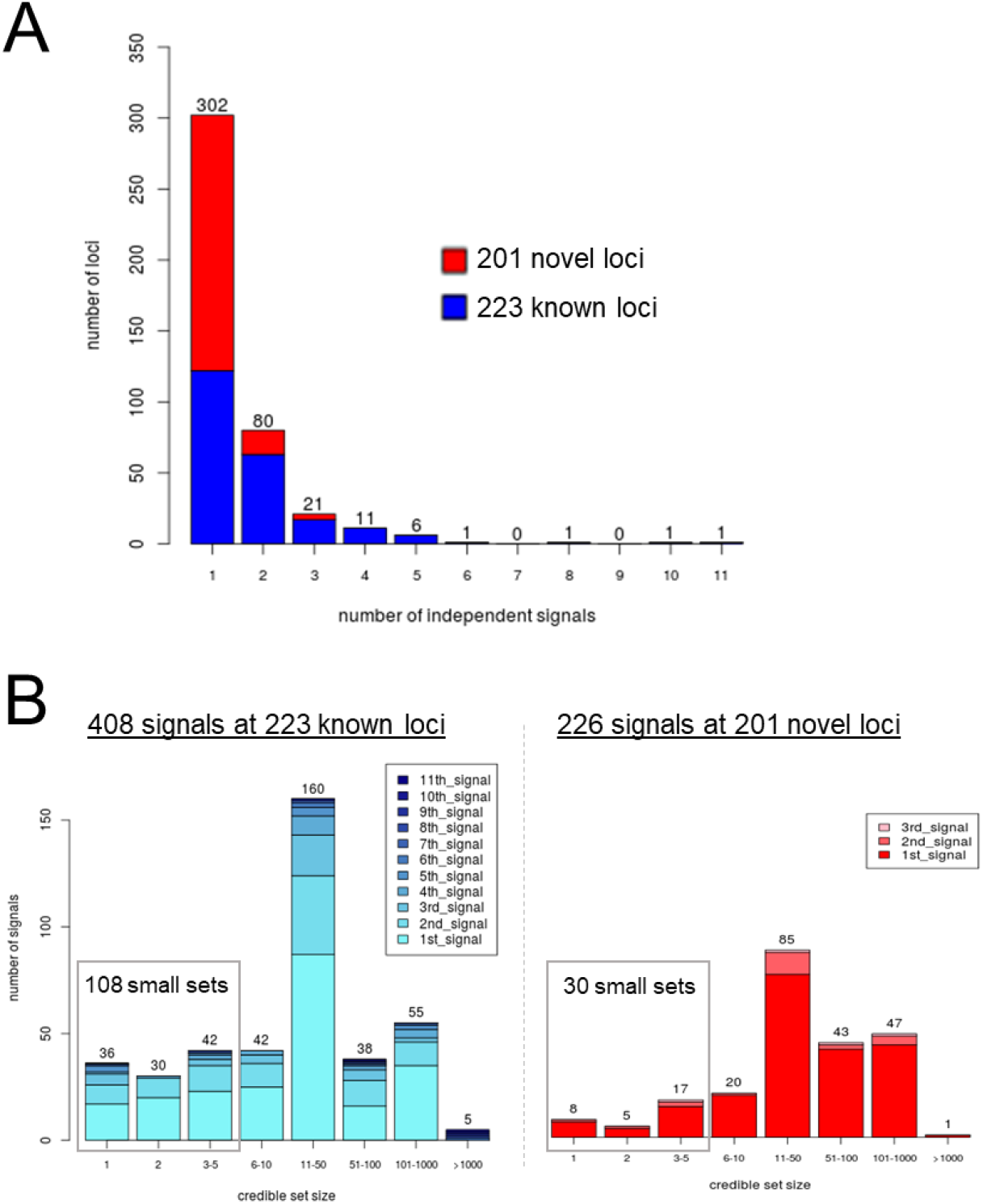
Distribution of independent signals and credible set sizes. For the 424 identified eGFRcrea loci, based on the European-only meta-analysis results (n = 1,004,040), we derived 634 independent signals by approximate conditional analyses with GCTA ^19^ and 99% credible sets of causal variants by fine-mapping using the method by Wakefield ^20^. A: Distribution of number of signals per 424 loci. B: Distribution of credible set sizes separately for the 408 signals at known and the 226 signals at novel loci (color denotes the order in which the signal appeared in the stepwise conditional analysis).

For statistical fine-mapping, we calculated the posterior probability of association (PPA) ^20^ for each variant and 99% credible sets of variants at each of the 634 signals (**Supplementary Table 6, Methods**). We observed 138 signals with small 99% credible sets (≤ 5 variants; 30 at novel and 108 at known loci) including 44 single-variant sets (8 at novel and 36 at known loci; **Figure 3B**, **Supplementary Table 7**). Overall, these include 30 single-variant sets that were not reported as single-variant sets previously by Wuttke *et al.* (8 at novel and 22 at known loci), either because the respective locus or signal was unidentified or because the credible set was larger. For signals in known loci, the increased GWAS sample size substantially reduced 99% credible set sizes compared to previous work ^7^ (median 99% credible set size = 17 variants versus 26 previously, **Supplementary Table 7**).

To understand the overlap of primary GWAS lead variants with credible sets of variants derived in the European-only analyses, we compared the PPA of the primary GWAS lead variant with the highest observed PPA within the respective (European-ancestry) credible set (**Supplementary Figure 6**). Most of the 424 primary lead variants were precisely the variant with the highest PPA (215 variants) or were contained (158 variants) in the respective European-ancestry credible set.

In summary, we identified 634 independent signals of which 138 (~22%) showed high fine-mapping resolution down to five or less 99% credible set variants, including 44 signals with only one variant. The signals with small credible sets might be the most practical and effective to follow in a search for the causal variants, as these contain most likely the causal variant – under the assumption that there was exactly one causal variant and that this variant was among those analyzed. We thus utilized the credible set variants to aid prioritizing likely causal variants and genes in the next step.

### Gene prioritization

In order to guide future functional follow-up of identified loci, we aimed to prioritize genes using bioinformatic analyses and so provide a searchable and sortable Gene PrioritiSation (GPS) table. To explore genes residing in or near the GWAS signals, we selected the 5,907 candidate genes overlapping the 424 locus regions (i.e. first and last GWS variant of a locus +/−250kb; 2,088 or 3,819 genes at novel or known loci, respectively **Supplementary Table 7**): the average number of genes per locus was 8. A maximum of 363 genes mapped to the ‘k73’ locus (MHC). We then screened the credible set variants and genes for the following GPS features: (i) variant with relevant predicted function based on Combined Annotation-Dependent Depletion (CADD) PHRED-like score ^21^ (focusing on variants within candidate genes, CADD-Score ≥15), (ii) variant with relevant regulatory function based on significant expression in tubulo-interstitial or glomerular tissue from NEPTUNE ^22^ (FDR < 5%) or expression/splicing effects in kidney or other tissues from GTEx ^23^ (FDR < 5%), or (iii) genes with kidney-related phenotypes in mice (Mouse Genome Informatics ^24^, MGI) or human (Online Mendelian Inheritance in Man ^25^, OMIM^®^, or Mendelian kidney disease in Groopman and colleagues ^26^, **Methods**). Among the 5,907 candidate genes, we found 2,722 genes that showed at least one relevant GPS feature (**Table 1, Supplementary Tables 8-16**). This illustrates that considering one feature at a time provides limited information and that gene prioritization requires an aggregation of multiple features.

**Table 1:** Summary of Gene PrioritiSation (GPS) results. We applied bioinformatic follow-up analyses to the 5,960 candidate genes mapping to the 424 loci derived by our primary meta-analysis for eGFRcrea (n = 1,201,929). The table shows an overview of bioinformatic analyses and number of genes with evidence for the particular analysis by 201 novel and 223 known loci. Detailed results for the individual GPS features can be found in **Supplementary Tables 8-16**.

| Category | N° of genes with evidence |  |
| --- | --- | --- |
|  | Novel loci | Known loci |
| Stop-gained/ stop-lost/ non-synonymous (CADD ≥ 15) | 53 | 88 |
| canonical-splice/ noncoding-change/ synonymous/ splice-site (CADD ≥ 15) | 4 | 7 |
| Other consequence (CADD ≥ 15) | 103 | 140 |
| eQTL in glomerulus (NEPTUNE) | 6 | 11 |
| eQTL in tubulo-interstitium (NEPTUNE) | 22 | 76 |
| eQTL in kidney tissue (GTEx) | 7 | 32 |
| eQTL in other tissue (GTEx) | 733 | 1,552 |
| sQTL in kidney tissue (GTEx) | 6 | 24 |
| sQTL in other tissue (GTEx) | 346 | 711 |
| Kidney phenotype in mouse (MGI) | 115 | 226 |
| Kidney phenotype in human (OMIM, Groopman et al. N Engl J Med. 2019) | 77 | 152 |
| N° tested genes = 5,907 |  |  |
| N° genes with any entry: 2,722 |  |  |

While the quality of the bioinformatic follow-up approaches has improved markedly during the last decade due to larger GWAS data narrowing down credible sets of variants, larger more differentiated tissue data for eQTL analyses, and better prediction of genetic function ^12^, it is still unclear how to weigh the different sources of evidence to prioritize genes and variants for functional follow-up. Also, such a prioritization may depend on specific research question and on the available functional model. Therefore, we generated a searchable, sortable, and customizable GPS table, where the weights for each piece of evidence can be adapted to the specific needs of the researcher (**Supplementary Table 17**).

Here, we illustrate the usage of the GPS table by three examples and how this can derive interesting ideas. In the <u>first example</u>, we queried the GPS table for new biological hypotheses obtained from a list of prioritized genes located at novel kidney function loci by genes with at least three relevant GPS features. We thus filtered the GPS table for loci that were novel and eGFRcys/BUN-validated, set equal weights for (i) genes containing credible set variants that were protein-altering (CADD ≥ 15, ‘stop-gained’, ‘stop-lost’ or ‘non-synonymous’), (ii) genes mapping to credible set variants that were regulatory in kidney tissue (eQTL or sQTL, FDR<5%), and (iii) genes with a reported kidney phenotype in mice or human, and down weight other GPS features. This resulted in 12 genes at novel loci with strong evidence (gene score ≥ 3, **Figure 4A, Supplementary Note 1**). Among those, 10 genes scored in the category of human kidney-related phenotype including 9 linked to Mendelian kidney diseases according to Groopman et al. ^26^ *(ACTN4, BCS1L, BMP4, COL18A1, EVC, HNF1A, LAMC2, NPHP3, and NPHS1,* **Supplementary Table 16**). For two of these nine genes, we had a good idea about the potentially causal variant *(HNF1A* and *NPHS1;* mapping to small credible sets with 4 and 2 variants, including non-synonymous variants, rs1800574 and rs3814995, respectively, **Supplementary Table 6-8**). This supports the idea of phenotype associations in the general population that map to genes known for rare mutations related to severe phenotypes.

**Figure 4.**
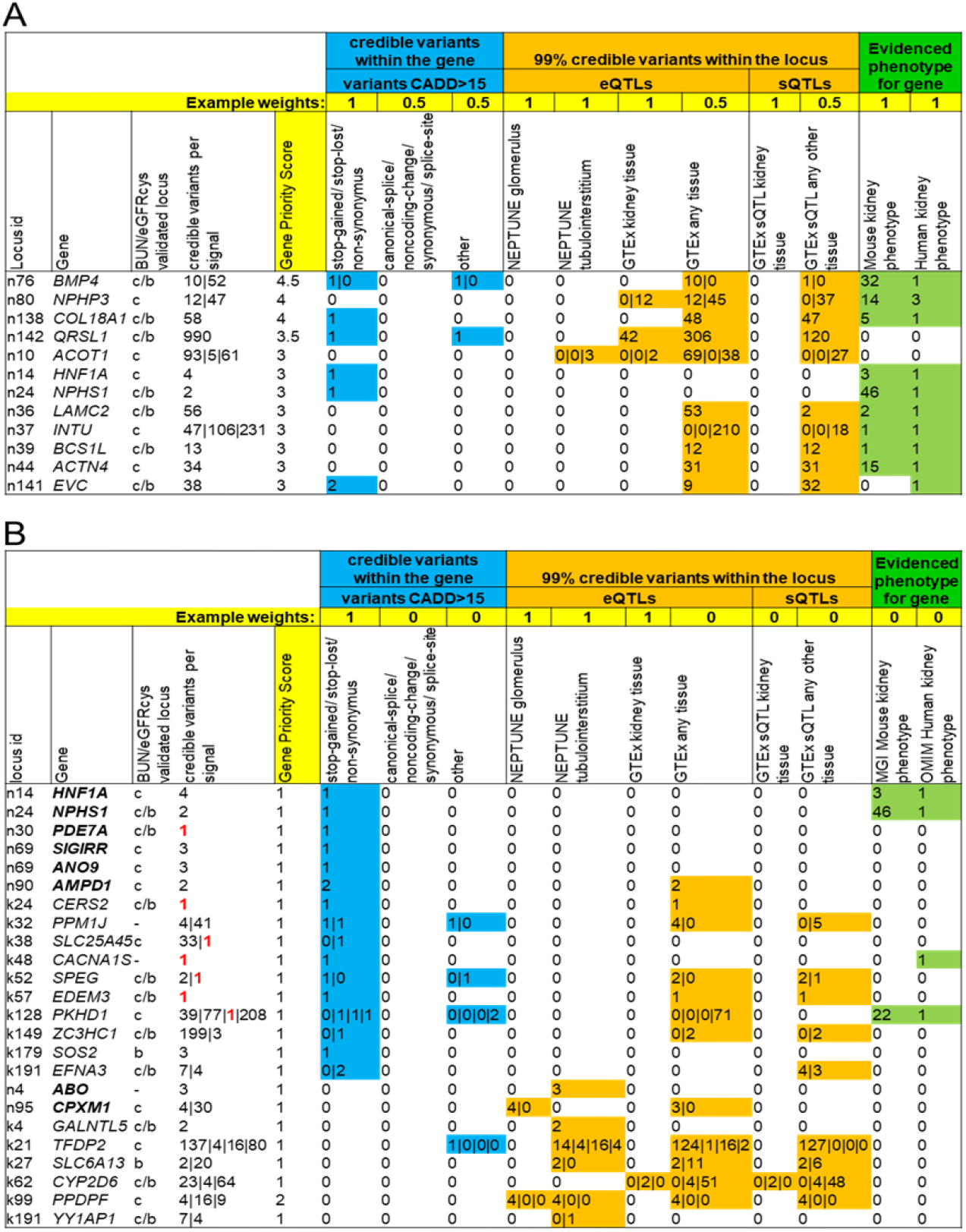
Gene prioritization results for 34 highlighted genes. The figure shows two example snippets of the GPS table (**Supplementary Table 17**). A: 12 highly-ranked genes (gene score >= 3) at novel loci that were found to be likely relevant for kidney function (i.e., validated by BUN or eGFRcys). B: 24 genes mapping to eGFRcrea signals with particularly small 99% credible sets (<= 5 likely causal variants) that contain protein-altering (‘stop-gained’, ‘stop-lost’ or ‘non-synonymous’) variants or eQTLs in kidney tissue. The snippets show whether the genes map to BUN (‘b) or eGFRcys (‘c’) validated loci, the number of 99% credible set variants per signal in the locus, the functional annotation of credible set variants within the gene (blue), regulatory evidence by credible set variants in the signal (orange) and whether the gene is known for kidney-related phenotypes in mouse or human (green). In panel B, genes mapping to novel loci are marked in bold and single-set variants are marked in red.

<u>In the second example</u>, we were interested in genes that mapped to small credible sets with protein-altering or kidney-tissue regulatory variants. We thus filtered the GPS table for small 99% credible sets (≤5 variants), set equal weights for genes containing credible set variants that were protein-altering (CADD ≥ 15, ‘stop-gained’, ‘stop-lost’ or ‘non-synonymous’) or eQTLs in kidney tissue (FDR<5%) and zero weights for other GPS features. This resulted in 24 genes with a relatively clear idea of what variant to follow as potentially causal variant (16 and 8 genes at known and novel loci, respectively, **Figure 4B**). Among the 16 genes in known loci, 10 contained a protein-altering variant (5 not indicated before ^7^: *EFNA3, PKHD1, SOS2, SPG* and *ZC3HC1;* 5 indicated before ^7^: *CACNA1S, CERS2, EDEM3, PPM1J* and *SLC25A45,* **Supplementary Table 8**) and 6 mapped to kidney-tissue eQTLs *(CYP2D6, GALNTL5, PPDPF, SLC6A13, TFDP2* and *YY1AP1,* **Supplementary Table 9-11**). Among the 8 genes in novel loci, 6 contained a protein-altering variant *(AMPD1, ANO9, HNF1A, NPHS1, PDE7A* and *SIGIRR,* **Supplementary Table 8**), the *CPXM1* mapped to an eQTL-variant in glomerulus (**Supplementary Table 9**) and the *ABO* to an eQTL-variant in tubulo-interstitium (**Supplementary Table 10**). All of the 8 genes except *ABO* were located in eGFRcys and/or BUN-validated loci. The 24 genes mapping to small credible sets included 6 genes mapping to a single protein-altering 99% credible variant (1 at a novel locus: *PDE7A;* 2 at known loci not indicated as single-set before ^7^: *CERS2* and *PKHD1;* and 3 indicated as single-set before ^7^: *CACNA1S, EDEM3,* and *SLC25A45).*

<u>In the third example</u>, when restricting to the 625 genes known for Mendelian kidney disease^26^, we found 212 among our candidate genes (33.9% of the 625) including four mapping to small ≤5 credible sets that contained protein-altering variants (novel loci: *NPHS1,* rs3814995; *HNF1A,* rs1800574; known loci: *PKHD1,* rs76572975; *CACNA1S,* rs3850625). All of these four genes were located in eGFRcys-and/or BUN-validated loci, except *CACNA1S,* which is known to be active in skeletal muscle cells ^27^ and thus unimportant for kidney disease.

Overall, the searchable and sortable GPS table and the three examples for GPS table reduction highlighted several interesting sets of genes: (i) 12 genes with interesting GPS features and located in novel loci that were eGFR- and/or BUN-validated, thus providing new hypotheses on potential kidney function genes, (ii) 24 genes with small credible sets containing a protein-altering or regulatory variant that provide an immediate idea for a potentially causal variant, and (iii) 212 genes among the known Mendelian kidney disease genes including four genes with small credible sets.

### Cell-type specific gene expression

Next, we were interested in the target cell types of the 5,907 candidate genes. We evaluated whether the genes were specifically expressed in relevant kidney cell-types using LDSC-SEG ^28^ (i.e., whether the gene was among the upper 10% of specifically expressed genes in the respective cell type**, Methods**). For LDSC-SEG analyses we used two independent single-cell RNA-seq data sets of human mature kidney ^29,30^. We found cell-type specific expression for multiple candidate genes (**Supplementary Table 18-19**). Of particular interest were two subsets of genes: (i) Among the 12 highly ranked genes located at novel kidney function loci (from **Figure 4A**), all except *QRSL1* were specifically expressed in at least one kidney cell type (**Table 2**). These include *BMP4* in loop of Henle, ascending vasa recta endothelium, glomerular endothelium, connecting tubule and principal cells, *NPHS1* in loop of Henle and podocytes. The *ACOT1* gene, for which we observed significant eQTLs in tubulo-interstitial tissue, displayed specific expression in proximal tubule in both data sets ^29,30^ (**Table 2**). (ii) Among the eight genes mapping to small credible sets (≤5 variants) with kidney-tissue eQTLs (from **Figure 4B**), all except *CYP2D6* were specifically expressed in at least one kidney cell type (**Table 2**). These included two genes at novel loci: *ABO* specifically expressed in connecting tubule, interstitial cells, loop of Henle, principal cells, proximal tubule, pelvic epithelium and transitional epithelium of ureter, and *CPXM1* in fibroblasts and podocytes.

**Table 2:**
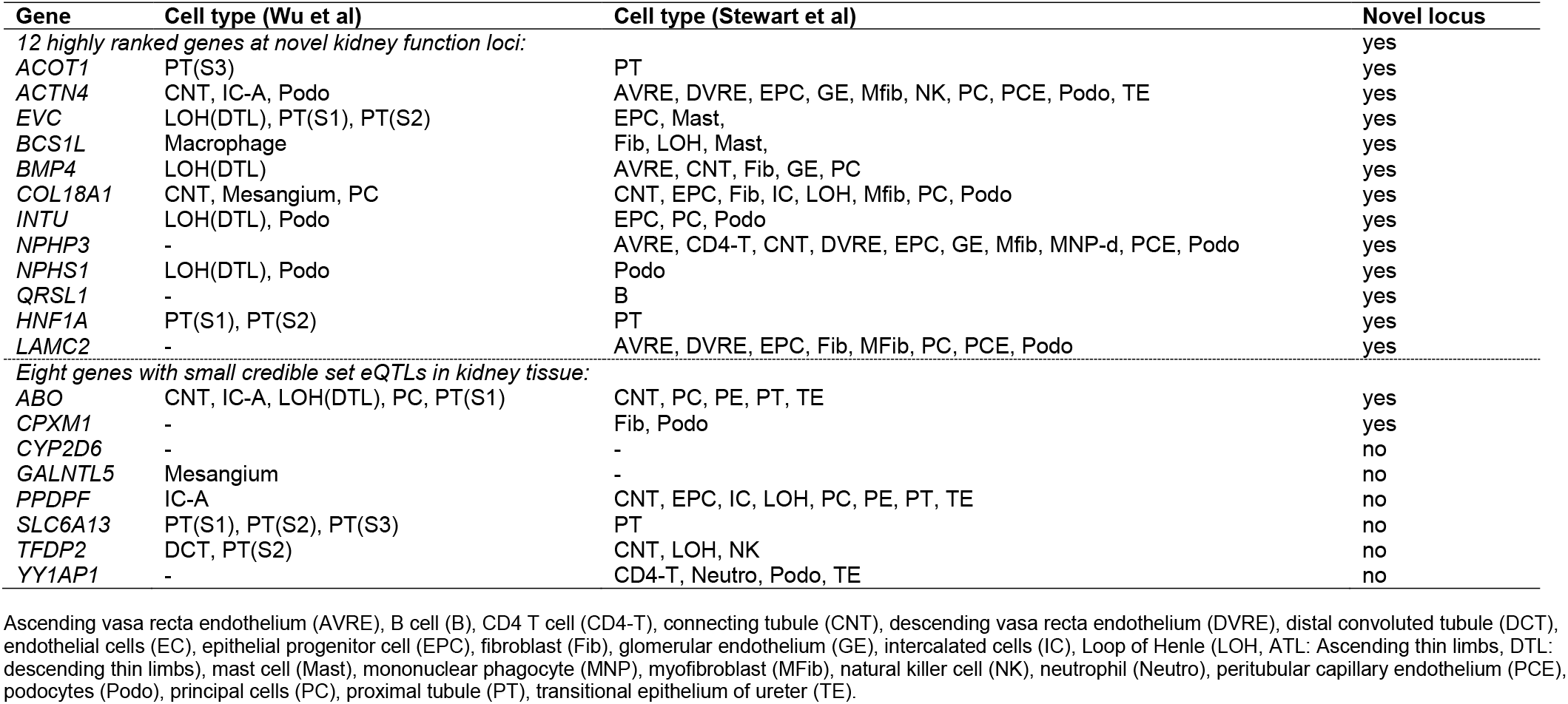
Cell-type-specific expression results for selected genes. The table summarizes cell-type specific expression results for the 12 highly ranked genes at novel kidney function loci (those from **Figure 4A**, **Supplementary Note 1**) and for the eight genes mapping to eGFRcrea signals with small 99% credible sets (<= 5 variants) that included significant eQTLs (FDR<5%) for the respective gene in kidney tissue (those from **Figure 4B**). The table lists cell types in which the respective gene is specifically expressed: Results for all genes are shown in **Supplementary Table 18** (expression data by Wu et al. ^30^) and **Supplementary Table 19** (expression data by Stewart et al. ^29^).

| Gene | Cell type (Wu et al) | Cell type (Stewart et al) | Novel locus |
| --- | --- | --- | --- |
| *12 highly ranked genes at novel kidney function loci:* |  |  | yes |
| ACOT1 | PT(S3) | PT | yes |
| ACTN4 | CNT, IC-A, Podo | AVRE, DVRE, EPC, GE, Mfib, NK, PC, PCE, Podo, TE | yes |
| EVC | LOH(DTL), PT(S1), PT(S2) | EPC, Mast, | yes |
| BCS1L | Macrophage | Fib, LOH, Mast, | yes |
| BMP4 | LOH(DTL) | AVRE, CNT, Fib, GE, PC | yes |
| COL18A1 | CNT, Mesangium, PC | CNT, EPC, Fib, IC, LOH, Mfib, PC, Podo | yes |
| INTU | LOH(DTL), Podo | EPC, PC, Podo | yes |
| NPHP3 | - | AVRE, CD4-T, CNT, DVRE, EPC, GE, Mfib, MNP-d, PCE, Podo | yes |
| NPHS1 | LOH(DTL), Podo | Podo | yes |
| QRSL1 | - | B | yes |
| HNF1A | PT(S1), PT(S2) | PT | yes |
| LAMC2 | - | AVRE, DVRE, EPC, Fib, MFib, PC, PCE, Podo | yes |
| *Eight genes with small credible set eQTLs in kidney tissue:* |  |  |  |
| ABO | CNT, IC-A, LOH(DTL), PC, PT(S1) | CNT, PC, PE, PT, TE | yes |
| CPXM1 | - | Fib, Podo | yes |
| CYP2D6 | - | - | no |
| GALNTL5 | Mesangium | - | no |
| PPDPF | IC-A | CNT, EPC, IC, LOH, PC, PE, PT, TE | no |
| SLC6A13 | PT(S1), PT(S2), PT(S3) | PT | no |
| TFDP2 | DCT, PT(S2) | CNT, LOH, NK | no |
| YY1AP1 | - | CD4-T, Neutro, Podo, TE | no |

Ascending vasa recta endothelium (AVRE), B cell (B), CD4 T cell (CD4-T), connecting tubule (CNT), descending vasa recta endothelium (DVRE), distal convoluted tubule (DCT), endothelial cells (EC), epithelial progenitor cell (EPC), fibroblast (Fib), glomerular endothelium (GE), intercalated cells (IC), Loop of Henle (LOH, ATL: Ascending thin limbs, DTL: descending thin limbs), mast cell (Mast), mononuclear phagocyte (MNP), myofibroblast (MFib), natural killer cell (NK), neutrophil (Neutro), peritubular capillary endothelium (PCE), podocytes (Podo), principal cells (PC), proximal tubule (PT), transitional epithelium of ureter (TE).

In summary, cell-type specific expression analyses provided further insights into potential target kidney cells highlighting numerous interesting biological candidates.

### Locus-based colocalization and a comparison with variant-based eQTL analysis

In the GPS, we analyzed the credible set variants, which were derived based on their association with eGFRcrea, for association with gene expression using a false discovery rate (FDR) approach. An alternative approach is the colocalization analysis comparing the eGFRcrea signal with the expression signal ^31^. We conducted colocalization analyses of eGFRcrea association and gene expression in tubulo-interstitial and glomerular tissue from NEPTUNE using ‘gtx’ (**Methods**). We found 55 and 18 colocalizations of eGFRcrea association signals with gene expression in tubule-interstitium or glomerulus, respectively (posterior probability of ‘positive’ colocalization, PP_H4_ ≥80%, **Supplementary Table 20-21**). The variant-based FDR approach and the signal-based colocalization usually provided similar results, particularly for small credible sets (≤5 variants, **Supplementary Figure 7**). For example, among the 8 genes that mapped to small credible sets with kidney-tissue eQTLs (from **Figure 4B**), four also showed a positive colocalization (PP_H4_≥80%) in the respective NEPTUNE kidney tissue *(GALNTL5, PPDPF, SLC6A13, YY1AP1* in tubulo-interstitium*; PPDPF* in glomerulus, **Table 3**). However, we also found examples for discordant results: for example, for *ABO*, we found an eQTL-variant in the small credible set, but no colocalization (PP_H4_<0.01 in tubule-interstitium), which may be attributed to unidentified independent signals in the variant-expression analysis (**Supplementary Figure 8**).

**Table 3:**
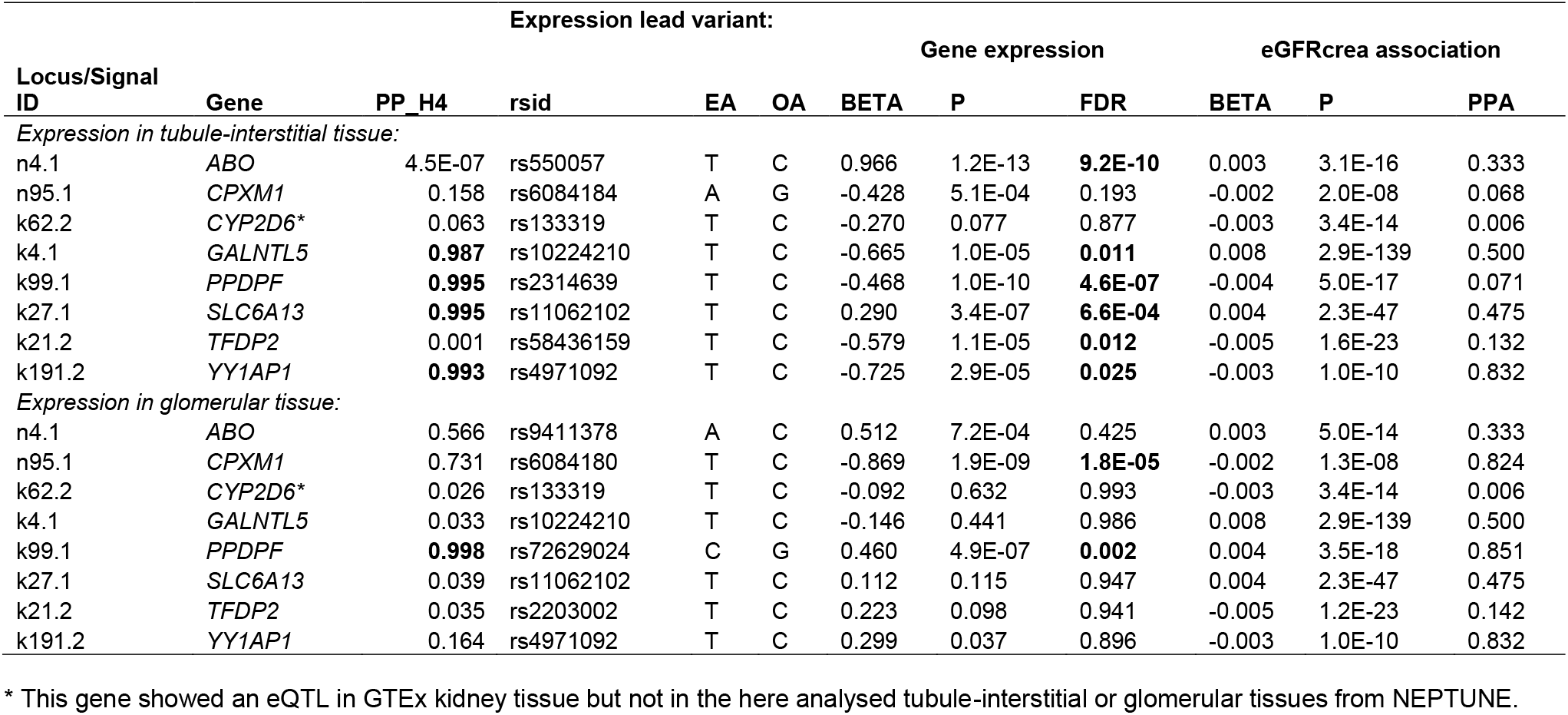
Colocalization analysis results for selected genes. For the eight genes with small 99% credible sets (<= 5 variants) that contain significant eQTLs in kidney tissue, the table shows results from colocalization analysis between eGFRcrea association and gene expression signals for two kidney tissues from NEPTUNE (tubule-interstitial and glomerular tissue, PP_H4 is the posterior probability of positive colocalization). The table also shows expression and eGFRcrea summary statistics for the credible variant with the smallest P-value for association on gene expression in the respective tissue (EA: Effect allele, OA: Other allele, BETA: genetic effect per EA, P: Association P-Value, FDR: False-discovery-rate, PPA: Posterior probability of association from variant-based fine-mapping). Locus/Signal ID: Identifier of identified locus/signal (‘n’ novel, ‘k’ known; first integer indicating the locus, second integer the signal within the locus). Marked in bold are positive colocalizations (PP_H4≥80%) and significant eQTLs (FDR<5%).

In summary, colocalization analyses show supportive results for many eQTL-findings among credible set variants in precisely the same kidney tissue, but not for all. The non-support can be due to chance, the possibly more limited power in the 2-stage approach inquiring eQTLs among credible set of variants, or possibly a colocalization signal that was more driven by the genetic eGFRcrea association than the expression evidence, which is based on a much smaller sample size.

### Aggregated genetic impact on eGFRcrea

To quantify the overall genetic impact on eGFRcrea, we estimated the genetic heritability and phenotypic variance explained, and derived a genetic risk score (GRS) effect based on eGFRcrea summary statistics (**Methods**). (1) Using LD-score regression ((LDSC))^32^, we estimated the additive contribution of all analyzed variants, i.e., narrow-sense heritability, h^2^=13.4% (based on unrelated individuals of European-ancestry from UKB, N = 364,674, **Table 4**). (2) LD-score regression analysis applied to specifically expressed genes (LDSC-SEG) ^28^ showed that eGFRcrea genetic heritability was significantly enriched (FDR<5%) in three proximal tubule clusters, principal cells and connecting tubule in expression data by Wu et al ^30^ and Stewart et al ^29^ (up to 2-fold enrichment, all novel findings except for two proximal tubule clusters ^33^, **Table 4, Supplementary Table 22**). The data from Stewart et al ^29^ were independent of the data from Wu et al ^30^ and were additionally analyzed here compared to the previous publication ^33^. (3) We estimated that 8.9% of the eGFRcrea variance was explained by the 634 independent signal lead variants (2.0% by the 226 novel signals’ lead variants and 6.9% by the 408 known ones, 7.6% by signals in eGFRcys/BUN-validated loci; based on European-only analysis; conditioned results in the case of multiple signals per locus; **Table 4, Supplementary Table 5**). (4) Finally, we observed significant and very similar GRS effects across two studies, in UKB as the largest source of our GWAS and in AugUR ^34^ as an independent population-based study (UKB, b_GRS_=-0.24 ml/min/1.73m^2^ per allele, P_GRS_ = 6.7 x 10^−5847^; AugUR, b_GRS_=-0.21 ml/min/1.73m^2^ per allele, P_GRS_ = 1.6 x 10^−9^; **Table 4**). In AugUR, individuals with a high versus low genetic risk (95% percentile of GRS versus 5%, i.e. 417 adverse risk alleles compared to 376) had on average −8.8 ml/min/1.73m^2^ lower eGFRcrea values.

**Table 4:**
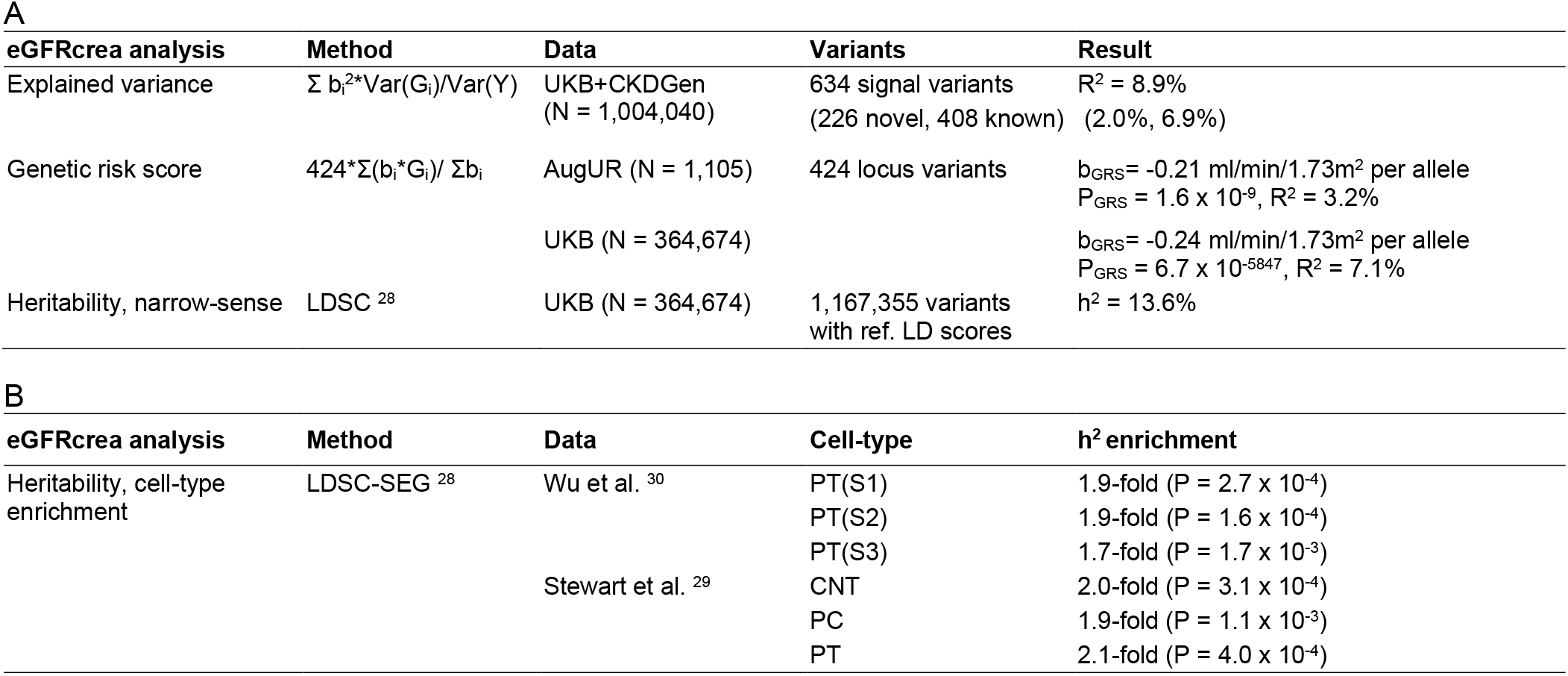
Aggregated genetic impact. The tables provide an overview of our analysis regarding the general genetic impact on eGFRcrea. A: Shown are results for explained variance, genetic risk score and heritability analyses for eGFRcrea. For each analysis, the table provides the method and data used as well as the main result. B: Shown are the significant heritability enrichments (FDR<5%) observed in various kidney cell-types (detailed results shown in **Supplementary Table 22**).

In summary, we found increased genetically explained eGFRcrea variability and enriched heritability in specific kidney cell-types and provided GRS estimates in an independent population-based study.

## DISCUSSION

In a meta-analysis of >1.2 million individuals from two of the largest GWAS data sets for eGFRcrea currently available, UKB and CKDGen, we identified 424 genomic loci for eGFRcrea, including 201 novel loci. Integration of eGFRcys and BUN in > 400.000 individuals supported 81% of the loci to be related to kidney function rather than creatinine metabolism. Across the 424 loci, we observed 634 independent variants that explained 8.9% of the eGFRcrea variance with the novel loci contributing a substantial fraction of ~2.0%. We documented a substantial impact of an adverse versus a beneficial genetic risk profile of almost −10 ml/min/1.73m^2^ in an independent population-based study. We aggregated comprehensive and systematic *in silico* follow-up results into a GPS tool to navigate through the abundance of evidence that provided several new hypotheses from novel loci and improved the fine-mapping resolution in known loci.

One challenge is the dissection of eGFRcrea loci for those being likely related to kidney function rather than creatinine metabolism. For this, we used genetic data on eGFRcys or BUN to assess consistency of effects. Kidney function assessment by eGFRcys is superior in predicting morbidity and mortality ^35^, but cystatin C measurement is expensive and less available in large epidemiological studies. BUN had been used previously to validate eGFRcrea GWAS results ^7^, but has known limitations ^36^. Furthermore, 75% of the eGFRcrea identified lead variants showed consistent association with eGFRcys compared to only 50% with BUN. Thus, eGFRcys might be more suitable to validate eGFRcrea association. Future work may improve the classification of kidney relevance by larger eGFRcys meta-analyses or by more complex clustering algorithms.

Selecting relevant genes for functional follow-up is also a challenging task in the interpretation of GWAS results. We here systematically aggregated *in silico* follow-up results into a GPS tool to help navigate through the abundance of evidence (**Supplementary Table 17**). The GPS is designed as a searchable and customizable tool to reflect different research questions and personal preferences, which should help guidance for functional follow-up studies. By various weighing and sorting examples, we illustrated the usage of this GPS and highlighted 34 interesting genes with novel insights.

We provide novel insights into known loci for eGFRcrea in various regards. Due to the increased sample size, we were able to substantially narrow down the association signal in known loci (**Figure 5A**): we found a median credible set size of 17 variants compared to 26 previously, 108 small credible sets compared to 58 and 36 single variant sets compared to 20. The small and particularly single variant sets are highly relevant when they include a variant with an interesting predicted function like protein-altering or kidney tissue regulatory. Our narrowing down of the signal will thus give more confidence into what variant and gene to follow for functional studies. For example, the *SOS2* gene maps to a credible set with 3 variants, including one protein-altering, compared to 9 variants in the credible set of this signal, previously ^7^. <u>Furthermore</u>, we successfully identified new independent signals within known loci: 185 secondary signals compared to 64 previously. When a locus is dissected into multiple signals, the credible sets per signal are usually more focused, which supports the narrowing down to the likely causal variants. For example, the dissecting of the locus containing the *PKHD1* gene into four independent signals compared to one signal previously enabled the new identification of a single variant credible set; this variant, rs76572975, is protein-altering with a MAF of 2% and located in the *PKHD1* gene. This gene is a known Mendelian disease gene ^26^ and the variant thus an immediately compelling target to explore further. Also, we identified four independent signals in the *UMOD/PDILT* locus compared to two signals previously, which indicates four independent functional entities that may be biologically interesting. This also led to a smaller credible set size for the second signal: 4 variants compared to 16 previously; of note the first signal was already a single variant set previously that we confirmed here.

**Figure 5.**
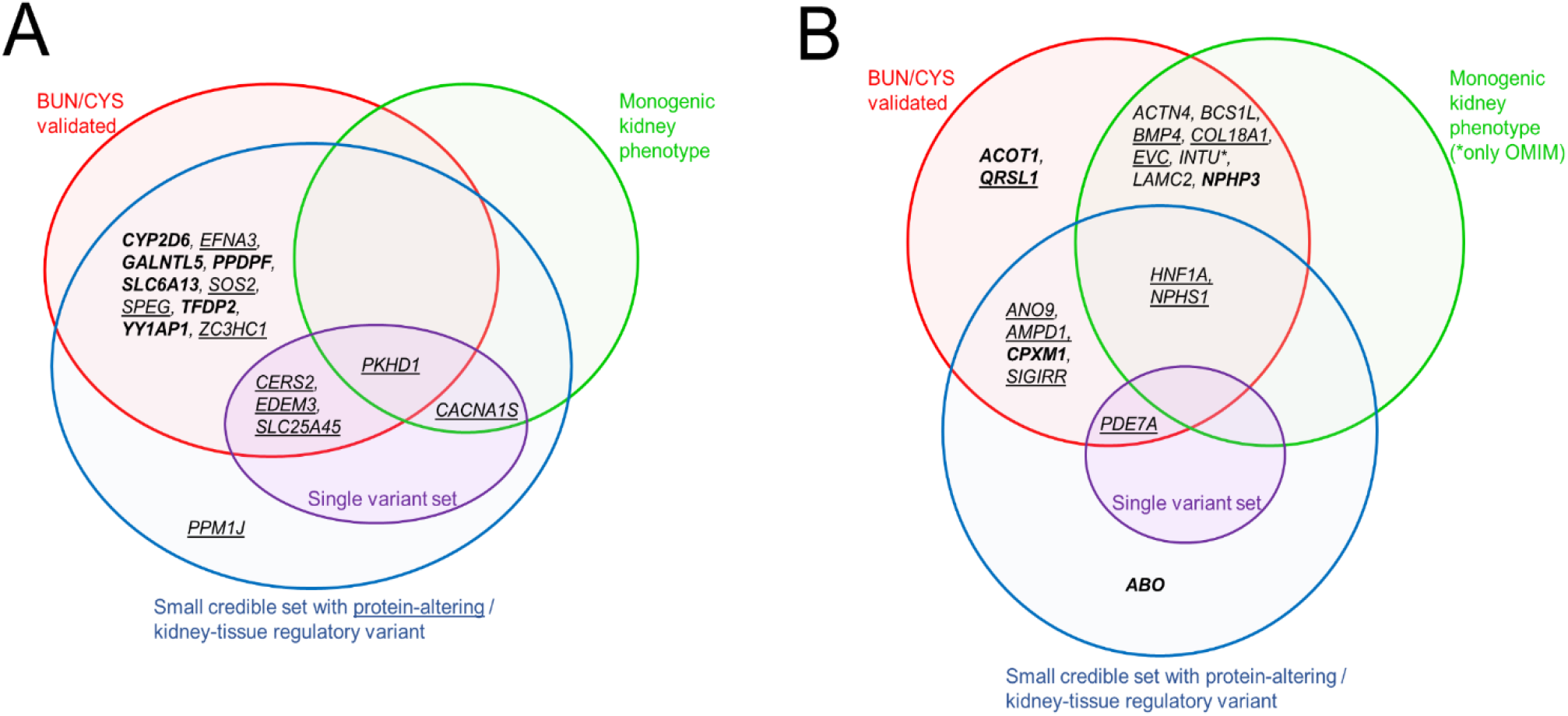
Overlap of gene prioritization evidence for 34 highlighted genes. The Venn diagrams show an overview on the interesting features of the 34 highlighted genes. A: 16 genes at <u>known loci</u> from **Figure 4B** (i.e., genes that mapped to small credible sets, ≤5 variants, containing protein-altering variants or kidney-tissue eQTLs). B: 18 genes at <u>novel loci</u> from **Figure 4A** (i.e., genes located at BUN/eGFRcys validated loci and showing a high gene score ≥3) or from **Figure 4B** (i.e., genes that mapped to small credible sets, ≤5 variants, containing protein-altering variants or kidney-tissue eQTLs). Genes mapping to protein-altering variants among the credible set are underlined, those mapping to kidney-tissue eQTLs are bold.

The novel loci and our GPS allowed us to highlight numerous genes with immediate ideas for a potentially causal variant and potentially novel biological hypotheses. We found nine Mendelian kidney disease genes ^26^ underneath nine novel loci with additional supporting features (**Figure 5B**). Each of these nine locus associations were validated by eGFRcys/BUN. The nine genes include *HNF1A* and *NPHS1,* which mapped to less-frequent or common protein-altering variants in small credible sets. One may speculate that these are variants with a kidney phenotype in the general population that is linked to the rare Mendelian disorder, diabetes MODY type 3 ^37^ or Nephrotic syndrome type 1 ^38^, respectively. Another gene with compelling evidence is *PDE7A,* which maps to 99% credible set variant that consists of a single protein altering variant, rs11557049. This variant is also the locus lead variant and showed consistent eGFRcys/BUN association. This makes this gene - that has not been reported before for kidney function – a likely kidney function gene with an immediate idea of a potentially causal variant to follow in functional studies. Further genes at novel loci with small credible sets containing protein-altering or kidney tissue regulatory variants are *ANO9, AMPD1, CPXM1, SIGIRR* all in eGFRcys/BUN-confirmed loci. These genes might be interesting to evaluate in whole exome sequencing data from patients with kidney disease of unknown origin.

The absence of independent replication for the 424 identified lead variants might be considered a limitation. However, the necessity of replication of GWAS findings has recently been revisited in the light of the general lack of suitable and appropriately powered replication samples ^39^. Still, all 424 lead variants showed directionally consistent nominal significant associations in UKB and CKDGen when analyzed separately (**Supplementary Table 1**). This supports our confidence into these associations being genuine. Another limitation is the lack of reference data for linkage disequilibrium in non-European ancestry individuals, which had us limit our fine-mapping on European-ancestry. Future studies augmenting on non-European ancestry individuals are warranted to provide equally powered analysis and fine-mapping for non-European ethnicities ^40,41^. Finally, our analyses were limited to Single Nucleotide Polymorphisms (SNPs) and disregarded structural variations, insertions or deletions. The reason for this was that the CKDGen summary statistics included only SNPs ^7^ and we thus restricted our meta-analysis accordingly.

In summary, our results help guide future functional follow-up studies on various ends: (i) the novel identified loci may generate new biological hypotheses, (ii) the improved fine-mapping resolution in the known loci will help select promising targets, and (iii) our searchable and customizable GPS table provides a powerful tool to support the cross-talk between GWAS researchers and molecular biology scientists.

## METHODS

### Study overview

We included two sources of GWAS data for eGFRcrea in our primary meta-analysis (n = 1,201,909): (i) GWAS summary statistics from the CKDGen consortium (n = 765,348, trans-ethnic) ^7^ that were downloaded from https://CKDGen.imbi.uni-freiburg.de and (ii) GWAS results generated in this work for eGFRcrea in UKB (application number 20272, n = 436,561, European ancestry) ^14^. We focused on European ancestry in UKB because this was the by-far largest ethnicity subset of UKB with other non-European ethnicities being clearly underrepresented and diverse ^14^. Secondary analyses included eGFRcrea meta-analyses of CKDGen and UKB based on individuals of Europeans ancestry only as well as meta-analyses of UKB and CKDGen for eGFRcys and BUN. Details on the phenotypes, downloaded data, association analyses, quality control, meta-analyses and further follow-up analyses are described in detail in the following.

### Phenotypes

The primary outcome of our meta-analysis is log-transformed eGFRcrea. This was used by the studies contributing to the CKDGen meta-analyses and for the UKB association analysis. In UKB, creatinine was measured in serum by enzymatic analysis on a Beckman Coulter AU5800 (UKB data field 30700, http://biobank.ctsu.ox.ac.uk/crystal/field.cgi?id=30700) and GFR was estimated using the Chronic Kidney Disease Epidemiology Collaboration (CKD-EPI) formula ^42,43^. For all studies involved in the CKDGen analysis, creatinine concentrations were measured in serum and GFR was estimated based on the CKD-EPI (for individuals > 18 years of age) ^42,43^ or the Schwartz (for individuals <= 18 years of age) ^44^ formula. Details on the study-specific measurements for the CKDGen studies were described previously ^7^. For all studies, eGFRcrea was winsorized at 15 or 200 ml/min/1.73m^2^ and winsorized eGFRcrea values were log-transformed using a natural logarithm. Secondary outcomes used for downstream analyses include log-transformed eGFRcys and log-transformed BUN. In UKB, cystatin C was measured based on latex enhanced immunoturbidimetric analysis on a Siemens ADVIA 1800 (UKB data field 30720, http://biobank.ctsu.ox.ac.uk/crystal/field.cgi?id=30720) and blood urea was measured by GLDH, kinetic analysis on a Beckman Coulter AU5800 (UKB data field 30670, http://biobank.ctsu.ox.ac.uk/crystal/field.cgi?id=30670). Details on the cystatin C and blood urea measurements in CKDGen studies can be found in previous work ^7,10^. In CKDGen and UKB, respectively, eGFRcys was obtained from cystatin C measurements using the formula by Stevens et al. ^45^ or the CKD-EPI formula ^42,43^. In all studies, eGFRcys was winsorized at 15 or 200 ml/min/1.73m^2^ and winsorized eGFRcys values were log-transformed using a natural logarithm. Blood urea measurements in mmol/L were multiplied by 2.8 to obtain BUN values in mg/dL, which were then log transformed using a natural logarithm.

### GWAS data from CKDGen

Each study in CKDGen had conducted GWAS for eGFRcrea while adjkusting for age, sex and other study-specific covariates. Summary statistics of each study were GC corrected. Details on study-specific analysis are described elsewhere ^7^. For our primary meta-analysis, we downloaded GWAS summary statistics for eGFRcrea from a trans-ethnic meta-analysis from the CKDGen consortium (https://CKDGen.imbi.uni-freiburg.de/files/Wuttke2019/20171016_MW_eGFR_overall_ALL_nstud61.dbgap.txt.gz, n = 765,348) ^7^. In this study, the authors had conducted trans-ethnic meta-analysis based on 121 GWAS results comprising 567,460 Europeans, 165,726 East Asians, 13,842 African-Americans, 13,359 South-Asians and 4,961 Hispanics. For our secondary downstream analyses, we also downloaded GWAS summary statistics for eGFRcrea from a European-only meta-analysis (https://CKDGen.imbi.uni-freiburg.de/files/Wuttke2019/20171017_MW_eGFR_overall_EA_nstud42.dbgap.txt.gz, n = 567,460) ^7^, for eGFRcys from a European-only meta-analysis (https://CKDGen.imbi.uni-freiburg.de/files/Gorski2017/CKDGen_1000Genomes_DiscoveryMeta_eGFRcys_overall.csv.gz, n = 24,061) ^10^ and for BUN from a trans-ethnic meta-analysis (https://CKDGen.imbi.uni-freiburg.de/files/Wuttke2019/BUN_overall_ALL_YL_20171017_METAL1_nstud_33.dbgap.txt.gz, n = 416,178) ^7^. Most studies included in the CKDGen meta-analyses were population-based. All studies used an additive genotype model and imputed the genotyped variant panel to the Haplotype Reference Consortium (HRC, v1.1) ^16^ or the 1000 Genomes Project (ALL panel) ^18^ reference panels. Details on the meta-analysis methods were described previously ^7,10^.

### GWAS data from UK Biobank

We conducted linear mixed model GWAS for log(eGFRcrea), log(eGFRcys) and log(BUN) in UKB using the fastGWA tool ^15^. We included age, age^2^, sex, age x sex, age^2^ x sex, and 20 principal components as covariates in the association analyses as recommended by the developers ^15^. The UKB GWAS were based on additively modeled genotypes that were imputed to HRC ^16^ and the UK10K haplotype reference panels ^17^. Details on the UKB genotypic resource are described elsewhere ^14^. We included individuals of European ancestry, i.e. self-reported their ethnic background as ‘White’, ‘British’, ‘Irish’ or ‘Any other white background’ (UKB data field 21000, http://biobank.ctsu.ox.ac.uk/crystal/field.cgi?id=21000). The sample sizes of the UKB GWAS were n = 436,581 for eGFRcrea, n = 436,765 for eGFRcys and n = 436,500 for BUN.

### Quality control

Prior to the meta-analysis, we applied a quality control (QC) procedure to the UKB and CKDGen GWAS results using EasyQC ^46^. We utilized the ‘CREATECPAID’ function to create unique variant identifiers that consisted of chromosomal, base position (hg19) and allele codes (i.e. ‘cpaid’, e.g. “3:12345:A_C”, allele codes in ASCII ascending order). For UKB, we excluded variants with a low imputation quality (Info<0.6) as done in the previous CKDGen analyses ^7^. For both data sets, UKB and CKDGen, we further excluded low frequency variants (MAF<0.1%) and any variants that were exclusively available in only one of the two data sets in order to limit analyses to variants that are available in UKB and CKDGen. This led to the exclusion of insertions, deletions and structural variants from the UKB GWAS results, since CKDGen focused on SNPs ^7^. We corrected our UKB association statistics for population stratification using the genomic control inflation factor (λ = 1.41) ^47^. We also calculated the genomic control inflation factor for the CKDGen results (λ = 1.32) but did not apply the correction because the individual studies contributing to the CKDGen meta-analyses were already GC corrected (see ^7^ for details on the study specific methods).

### Meta-analyses

We conducted fixed-effect inverse-variance weighted meta-analyses of CKDGen and UKB association results using metal ^48^. As primary meta-analysis, we combined log(eGFRcrea) association results from CKDGen (trans-ethnic) and UKB (n = 1,201,909). After meta-analsyis, we excluded variants with a low minor allele count (MAC < 400) yielding 13,633,840 variants in our final meta-analysis GWAS result for eGFRcrea. The GC lambda inflation factor of the eGFRcrea meta-analysis results was λ = 1.28 and the LD score regression intercept ^32^ was 0.90, which reflects conservative study-specific GC correction and indicates absence of confounding by population stratification. For downstream follow-up analyses, we also combined CKDGen European-only and UKB for log(eGFRcrea), as well as CKDGen and UKB for log(eGFRcys) and log(BUN).

### Locus definition and variant selection

We defined locus borders by adding +/− 250kb to the first and last GWS variant of a specific region. To achieve independent loci, we selected the variant with the smallest association P-value genome-wide as a starting point and defined this variant as the lead variant for its locus. Starting at the outermost two GWS variants (P < 5×10^−8^) in a 1Mb region centered on the lead variant, areas of another 500kb were checked for GWS variants. If GWS variants were found in this extended region, the region extension step was repeated on the novel outermost GWS variants until no further GWS variants were found. The positions of the two, last found GWS variant in both directions minus/plus 250kb were defined as the locus limits. The locus variants were omitted from the data and the whole process was repeated until no GWS variants remained genome-wide. We defined a locus as novel when none of the 264 known loci discovered by Wuttke et. al. ^7^ overlapped with our GWS variants. We used the so-defined locus regions (GWS variants +/−250kb cis window) for the *in silico* follow-up analyses and defined the genes that overlapped this locus regions as candidate genes.

### Validation for kidney function based on eGFRcys and BUN

To evaluate the eGFRcrea-associated lead variants for their potential relevance for kidney function, we analyzed their genetic association with log(eGFRcys) and log(BUN). Consistency of the eGFRcrea association for a given effect allele with eGFRcys- or BUN-association was defined as nominal significant association (P<0.05) and concordant effect direction for eGFRcys or opposite effect direction for BUN, respectively.

### Approximate conditional analyses using GCTA

To identify independent secondary signals at the identified loci, we conducted approximate conditional analyses based on European-only meta-analysis summary statistics using GCTA ^19^. The analysis was limited to European-only results due to the lack of an appropriate LD reference panel that mirrors the ethnicities in our primary meta-analysis of CKDGen (trans-ethnic) and UKB. We thus created a LD reference panel based on 20K randomly selected unrelated Europeans from UKB. For each identified locus, we applied a stepwise approach to derive the further signals: (i) we first conditioned on the locus lead variant and then selected the most significant variant across all locus variants in this conditional analysis. (ii) If this selected variant showed a genome-wide significant conditional P value (P_Cond_ < 5.0×10^−8^), this variant was deemed as independent signal lead variant and added to the list of variants to condition on. (iii) The procedure was repeated until no more genome-wide significant variant was identified.

### Credible sets of variants

For each variant in each of the identified signals, we calculated approximate Bayes factor (ABF) and PPA using the Wakefield method ^20^. We obtained 99% credible variant sets for each independent association signal. We used W = 0.005^2^ as prior variance as done previously ^7^. PPAs were calculated based on the meta-analysis summary statistics for loci with only one signal and based on conditioned summary statistics for loci with multiple independent signals (each signal conditioning on the other signal lead variants in the locus). One set of 99% credible variants was obtained for each independent signal.

### Gene PrioritiSation

To prioritize genes among the list of candidate genes at the discovered eGFRcrea loci, we performed a series of statistical and bio-informatic follow-up analyses based on the secondary signal analysis from the EUR-only meta-analysis. (1) For each credible set variant within a candidate gene we derived the CADD PHRED-Score ^21^ to prioritize credible variants concerning their potential protein altering outcome (CADD >= 15). We chose the threshold of 15, since this is representing the 3.2% most deleterious variants of the 8.6 billion variants available in CADD. CADD uses the Ensembl Variant Effect Predictor (VEP) ^49^ to obtain gene model annotation and combines this information to 17 possible consequence levels. Based on the CADD internal consequence score (ConsScore), we classified each prioritized variant into three groups: (i) ‘stop-gained’, ‘stop-lost’, ‘non-synonymous’ (ConsScore 8 or 7), (ii) ‘canonical-splice’, ‘noncoding-change’, ‘synonymous’, ‘splice-site’ (ConsScore 6 or 5), and (iii) other (ConsScore 4-0). We restricted to variants located within candidate genes to increase the percentage of variants with predicted protein altering outcome and to avoid major overlap with variants that influence gene expression levels, that are analyzed in the next steps.(2) We highlighted credible set variants within each locus that are significant expression quantitative trait loci (eQTLs) in kidney (and other) tissue for the related candidate genes. Therefore, we analyzed eQTLs quantified from glomerular and tubule-intestitial tissue in the NEPTUNE study ^22^ and from 44 tissues including kidney cortex in the GTEx project ^23^ with regard tosignificant association (FDR <0.05) on candidate gene expression levels. We used the FDR provided by GTEx and applied a Benjamini-Hochberg FDR correction ^50^ to the NEPTUNE association P-values for glomerular and tubule-intestitial tissue separately (to obtain a FDR for each variant x gene combination). We cut back on credible variants to link the information of the statistical association with eGFRcrea to the influence on the candidate gene expression.(3) To identify credible variants with significant effect (FDR <0.05) on expression levels of exon junctions or variation in the relative abundances of gene transcript isoforms (sQTLs) we used sQTL summary statistic from the GTEx database ^23^. (4) Kidney-relevant phenotypes in mice were selected from the Mouse Genome Informatics (MGI) ^24^ hierarchical ontology. All phenotypes subordinate to “abnormal kidney morphology” (MP:0002135) and “abnormal kidney physiology” (MP:0002136) were gradually extracted. A table with all genes occurring in MGI-database and the associated phenotypes was restricted to the kidney-relevant phenotypes and compared to the list of candidate genes. (6) We selected genes which are known to cause monogenic kidney phenotypes or disease in human based on two resources: the Online Mendelian Inheritance in Man (OMIM) ® database ^25^ and a recent publication by Groopman et al ^26^. We generated a table of kidney phenotypes and causal genes in the context of human disorders by querying the OMIM database for phenotype entries subordinate to the clinical synopsis class “kidney”. This table was manually reviewed and diseases with “kidney” - phenotype entries being: “Normal kidneys”, “Normal renal ultrasound at ages 4 and 7 (in 2 family)”, “No kidney disease”, “No renal disease; Normal renal function”, “Normal renal function; No kidney disease”, “No renal findings” were excluded. We further used a summary table by Groopman et al., which included 625 genes associated with Mendelian forms of kidney and genitourinary disease (http://www.columbiamedicine.org/divisions/gharavi/files/Kidney_Gene_List_625.xlsx). Both tables were combined and checked for concordance with candidate genes.

### Cell-type specific expression and LDSC-based heritability enrichment

We were interested in whether the candidate genes were specifically expressed in certain cell types. We used 17 human cell-types from Wu et al ^30^ and 27 human cell types from Stewart et al ^29^ and applied LDSC-SEG ^28^ analyses to obtain the top 10% specifically expressed candidate genes in each cell type. We queried our candidate genes and our highly prioritized genes (i.e., results from our gene prioritization) against the cell-type specific lists of genes. We were further interested in whether the genetic contribution to eGFRcrea differs between specific cell-types. We investigated whether heritability of eGFRcrea was enriched in one of the 17 or 27 kidney cell types from Wu et al ^30^ or Stewart et al ^29^. For each cell-type, we conducted LDSC ^32^ heritability analyses that were restricted to regions surrounding (+/−100kb to transcribed regions) the 10% most specifically expressed genes for the specific cell type. Details on the cell-type specific expression and heritability enrichment analyses and for the Wu et al data ^30^ can be found elsewhere ^33^. The Stewart et al data set including the expression matrix and cell-type annotation was downloaded from http://www.kidneycellatlas.org/ ^ref 29^.

### Colocalization analyses

We were interested whether our identified eGFRcrea signals co-localized with gens expression signals in tubule-interstitial or glomerular tissue from NEPTUNE ^22^. We conducted colocalization analyses using the method described by Giambartolomei et al ^31^. For each signal, colocalization analyses were performed for the respective locus’ genes separately for tubule-interstitial or glomerular tissue. We used the eGFRcrea EUR meta-analysis summary statistics for loci with only one signal and the conditioned summary statistics for loci with multiple independent signals. We used the R package ‘gtx’ and its coloc.compute function with 0.005^2^ as prior variance for the eGFRcrea association (similar to what was used for the statistical fine-mapping of credible variants) and 0.55^2^ as prior variance for the expression in tubule-interstitial or 0.53^2^ as prior variance for the expression in glomerular tissue. The prior variances for the expression data were obtained from the Wakefield formula (8) ^20^, assuming that 95% of significant eQTLs (FDR<0.05) in NEPTUNE fall within the effect size range −1.07 to 1.07 in tubule-interstitial or −1.04 to 1.04 in glomerular tissue.

### Heritability and explained variance

We were interested in the general impact of genetics on eGFRcrea. For this, we conducted various analyses including analyses of narrow-sense heritability (estimates the additive genetic contribution to eGFRcrea) and explained variance. We conducted LD score regression analyses using LDSC ^32^ to estimate narrow-sense heritability based on the UKB summary statistics for eGFRcrea (not GC-corrected, limiting to variants available in the LDSC reference data ‘w_hm3.snplist’). Explained variance was calculated for each of the independent signal lead variants and then summed up to obtain variance explained by all identified signal lead variants. For each variant, we calculated R^2^ = β^2^*Var(G)/Var(Y). Here, β is the genetic eGFRcrea effect (based on EUR meta-analysis summary statistics for loci with only one signal and conditioned EUR summary statistics for loci with multiple signals), Var(G) is the genetic variance calculated from Var(G)= 2*MAF*(1-MAF) and Var(Y) is the phenotypic variance (set to 0.016 as variance of age- and sex-adjusted log(eGFRcrea) residuals in 11,827 Europeans of the ARIC study as utilized previously ^7^).

### Genetic risk score analyses

To estimate an average and cumulative effect of genetic variants on eGFRcrea, we conducted GRS analyses in two studies: The German AugUR study (prospective study in the mobile elderly general population around Regensburg, Germany, age range 70-95y, mean +/− SD eGFRcrea = 70.0 +/− 15.5 ml/min/1.73m^2^, n = 1,105) ^34^, and UKB (prospective cohort study from UK, age range 40-69y, mean +/− SD eGFRcrea = 90.6 +/− 13.2 ml/min/1.73m^2^, n ~ 500,000) ^14^. While UKB was part of our variant identifying discovery meta-analysis, AugUR was an independent study. To obtain GRS effects that are interpretable as eGFRcrea units per allele, we did not apply a log-transformation to eGFRcrea for the GRS analyses. In both studies, we focused on unrelated individuals of European ancestry and calculated residuals of eGFRcrea from a linear model that adjusted eGFRcrea for age, sex and principal components (4 PCs for the AugUR and 10 PCs for UKB). We calculated a weighted and scaled GRS by adding up the eGFRcrea-decreasing alleles of the identified variants, weighing by the absolute genetic effect observed in the primary meta-analysis and then scaled the GRS to the interval [0, 2*m], with m being the number of identified lead variants, to make the GRS effect approximatively interpretable as average eGFRcrea decrease per allele.

## Data availability

Summary genetic association results for UKB and the meta-analysis of UKB and CKDGen for log(eGFRcrea), log(eGFRcys) and BUN can be downloaded from www.genepi-regensburg.de/ckd or from https://ckdgen.imbi.uni-freiburg.de/. The GPS table is also available from www.genepi-regensburg.de/ckd.

## ACKNOWLEDGMENTS

We thank the CKDGen Consortium and its Analyst Group for sharing infrastructure and for the fruitful discussion and feedback on the project. This research was conducted using the UKBB resource (application no. 20272). AugUR cohort recruiting and management was funded by the Federal Ministry of Education and Research (BMBF-01ER1206, BMBF-01ER1507, to I.M.H.) and by the German Research Foundation (DFG HE 3690/7-1, to I.M.H.). Genome-wide genotyping for AugUR was funded by the University of Regensburg for the Department of Genetic Epidemiology. The computational work supervised by I.M.H. was funded by the German Research Foundation (DFG) – Project-ID 387509280 – SFB 1350 (to I.M.H.). I.M.H. received funding from the Deutsche Forschungsgemeinschaft (DFG, German Research Foundation) – Project-ID 387509280 – SFB 1350 (subproject C6) and from the National Institutes of Health (NIH, R01RES511967). The work of A.K. was supported by DFG KO 2598/5-1. The DFG also supported this work within the funding programme Open Access Publishing.

## AUTHOR CONTRIBUTIONS

K.J. Stanzick, K.J. Stark, I.M.H. and T.W.W. wrote the manuscript; K.J. Stanzick, K.J.Stark, I.M.H. and T.W.W. conceived and designed the project; T.W.W. conducted association and genetic risk score analyses in UK Biobank, genetic risk score analyses in AugUR, approximate conditional analyses and colocalization analyses; K.J.Stanzick conducted meta-analyses, fine-mapping and gene prioritization analyses; Y.L. conducted cell-type specific expression analyses; A.K., M.G., M.W. and C.P. conceived and designed the CKDGen meta-analyses; All authors critically reviewed the manuscript.

