## Supplementary Figures for "Discovery and prioritization of variants and genes for kidney function in >1.2 million individuals"

#### Supplementary Figure 1. Analysis workflow.

General workflow of the analyses with indication of the sample sizes and data used. More details on the specific analyses are presented in the **Methods**. Abbreviations: MAF: minor allele frequency; Info: Imputation quality Info score; MAC: Minor allele count; eGFRcys: glomerular filtration rate estimated from cystatin, BUN: blood urea nitrogen, EUR: European, GCTA: Genome-wide Complex Trait Analysis; CADD: Combined Annotation Dependent Depletion; MGI: Mouse Genome Informatics; OMIM: Online Mendelian Inheritance of Men.

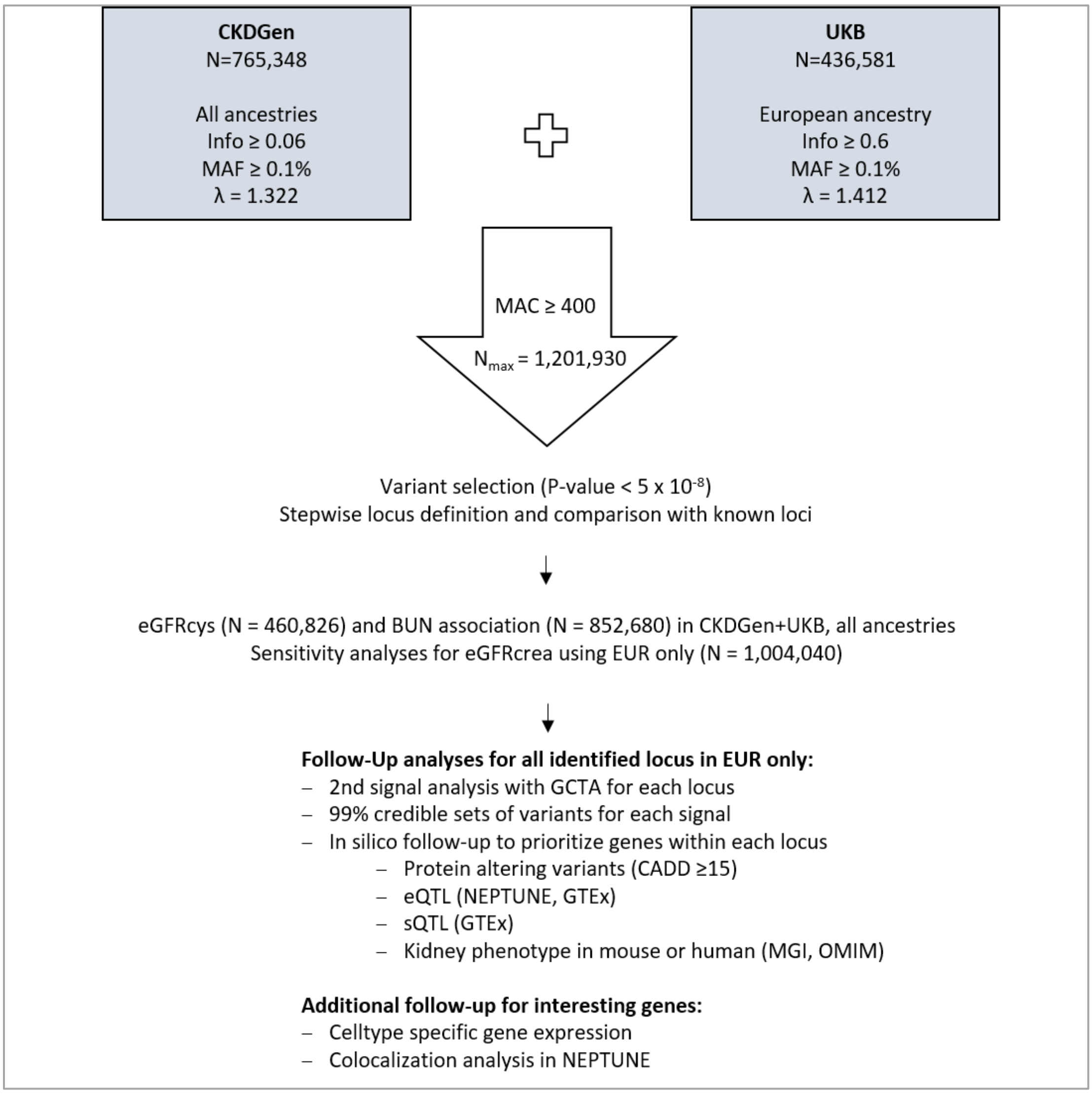

**Supplementary Figure 2. QQ-Plot for primary meta-analysis results for eGFRcrea.**

The QQ plot compares the expected versus the observed distribution of P-values in the primary meta-analysis for eGFRcrea (n = 1,201,929). Black dots mark association the $-\log_{10}$(P-values) of all variants; red dots show the respective results excluding the 264 loci previously described by Wuttke et al [1].

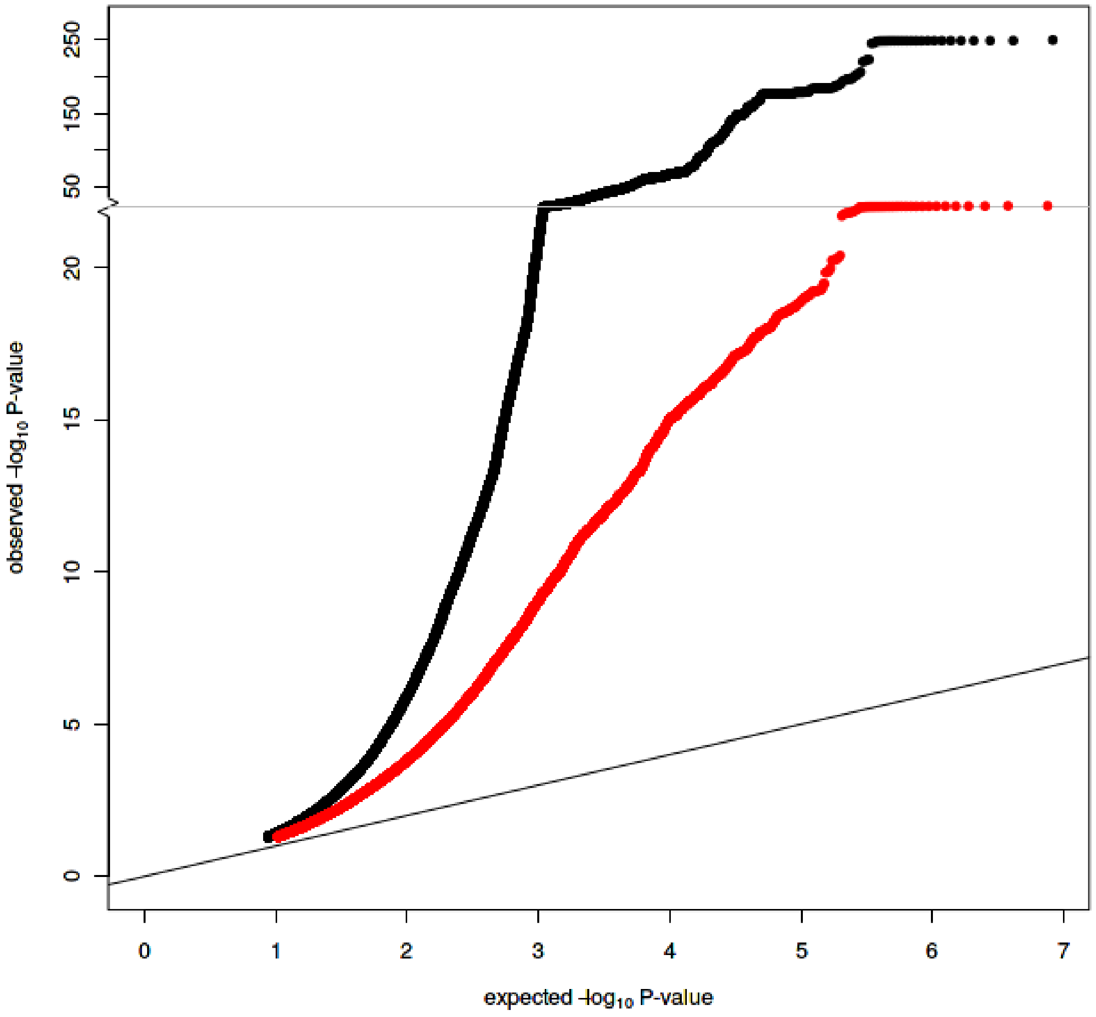

**Supplementary Figure 3. Comparison of primary and European-only meta-analysis for eGFRcrea.**

For the 424 lead variants identified by the primary meta-analysis (n = 1,201,929, including 197,888 non-European individuals), the scatterplots contrast the genetic effect sizes (aligned to eGFRcrea decreasing alleles) (**Panel A**) and the association P-Values (**Panel B**) between the primary and the European-only meta-analysis (n = 1,004,040).

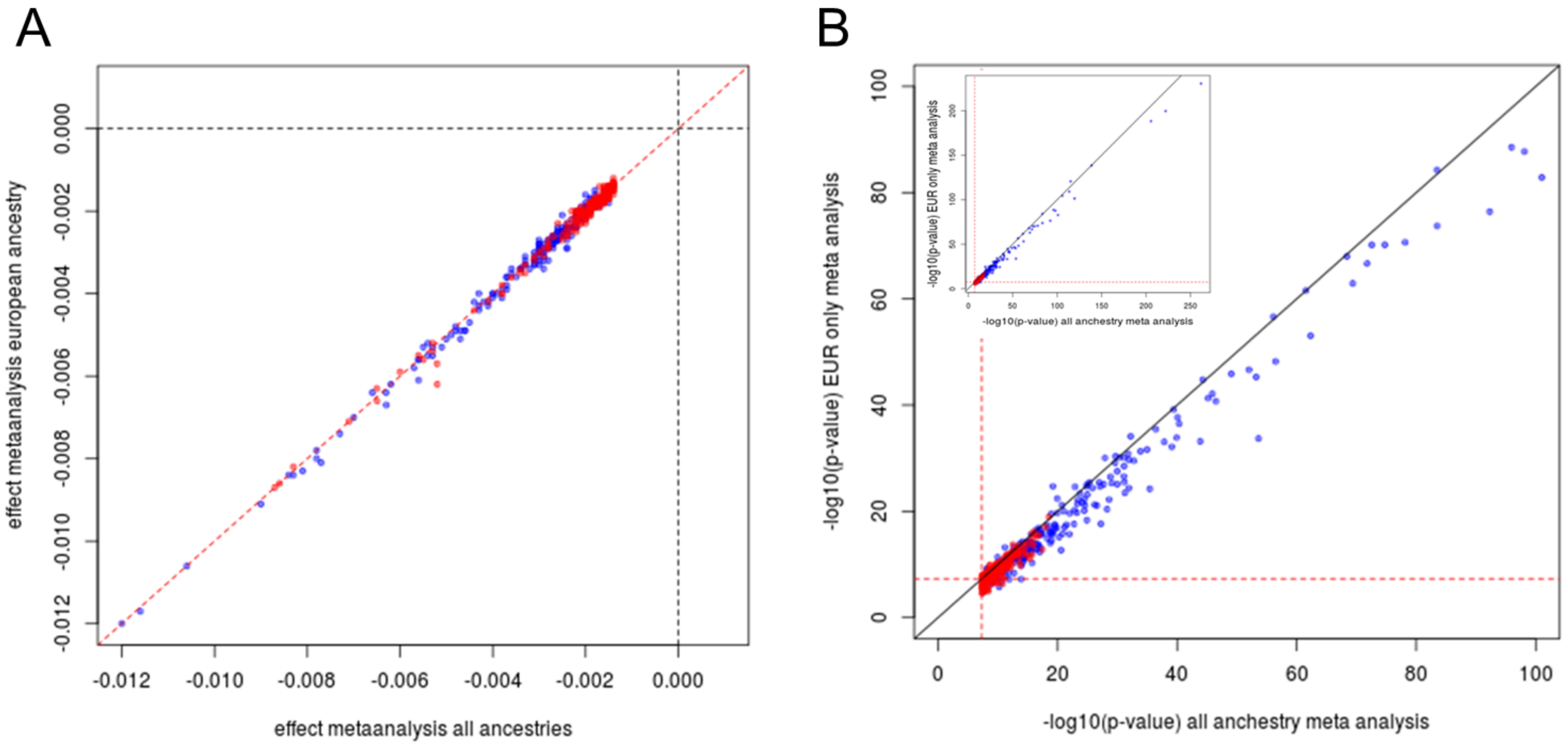

**Supplementary Figure 4. Regional Association Plots (RAPs) for eGFRcrea association with variants at the 21 novel loci that included multiple independent signals.**

The RAPs obtained with Locuszoom [2] show association results from the primary eGFRcrea meta-analysis (n = 1,201,929). Shown are 21 newly identified loci that included multiple independent signals in the stepwise approximate conditional analyses with GCTA [3]. P-values are reported without adjustment for the other signal(s) in the locus. Coloring denotes independent signals (using different colors) and correlation to the signal lead variants (from dark = highly correlated to bright = uncorrelated).

Locus n1:

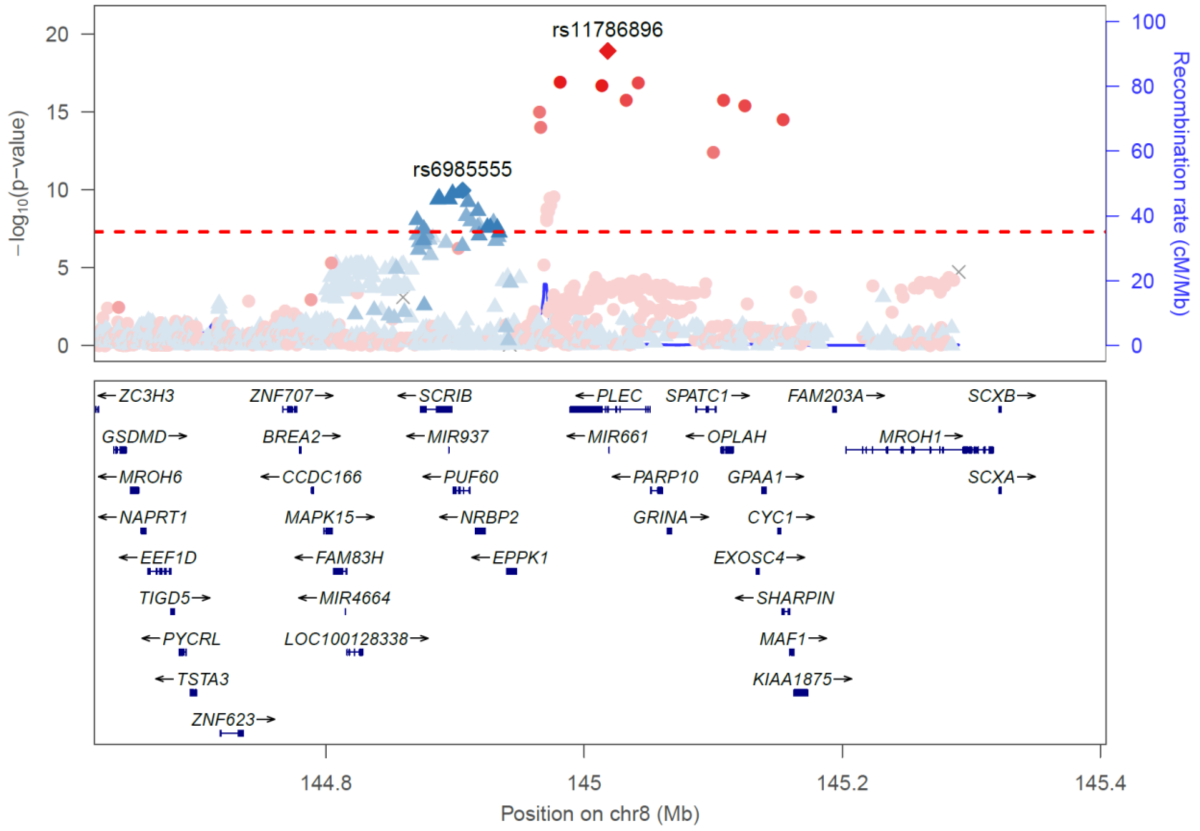

Locus n10:

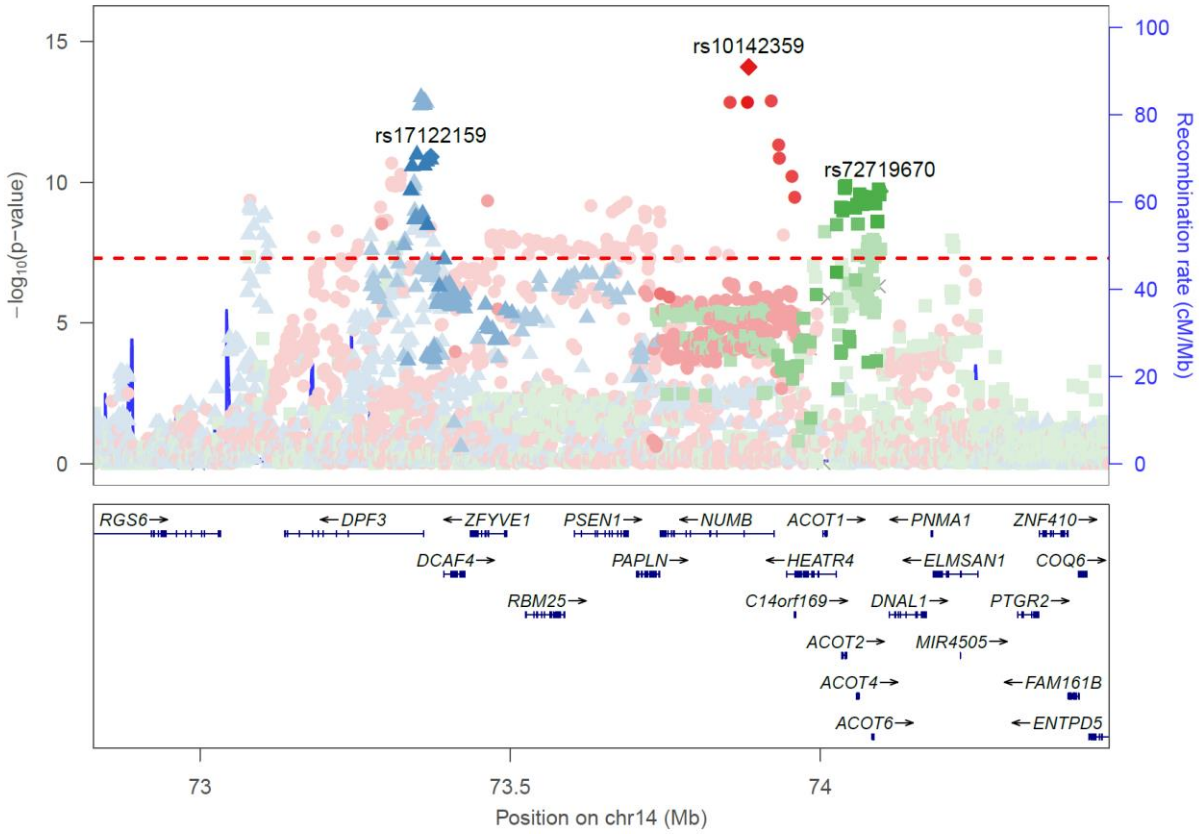


Locus n15

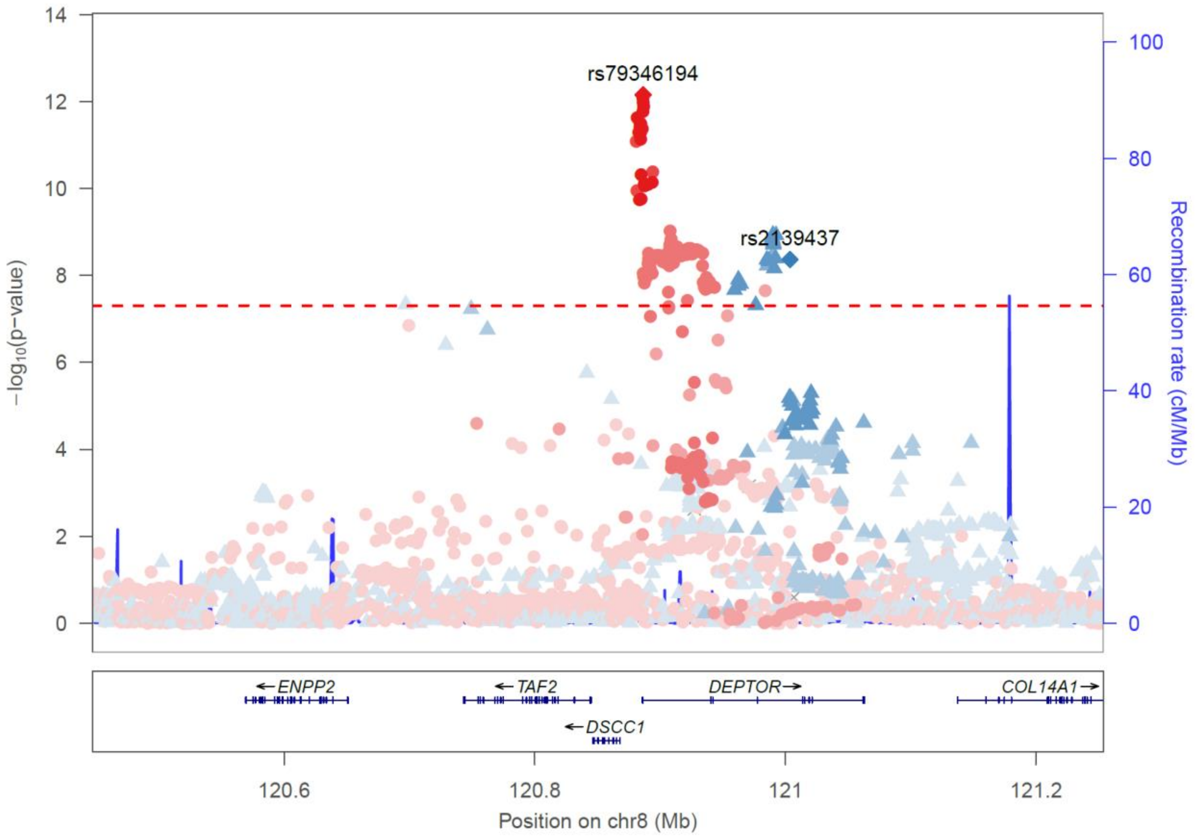

Locus n16

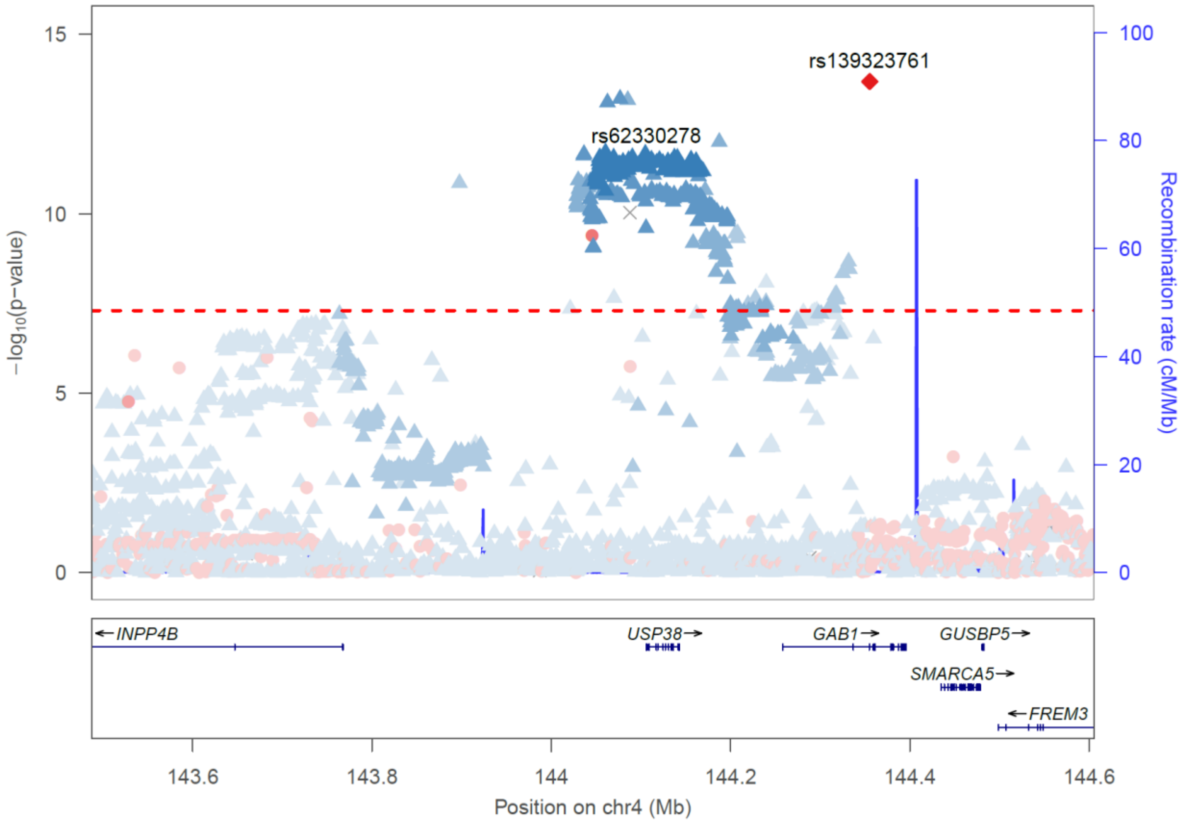


Locus n26

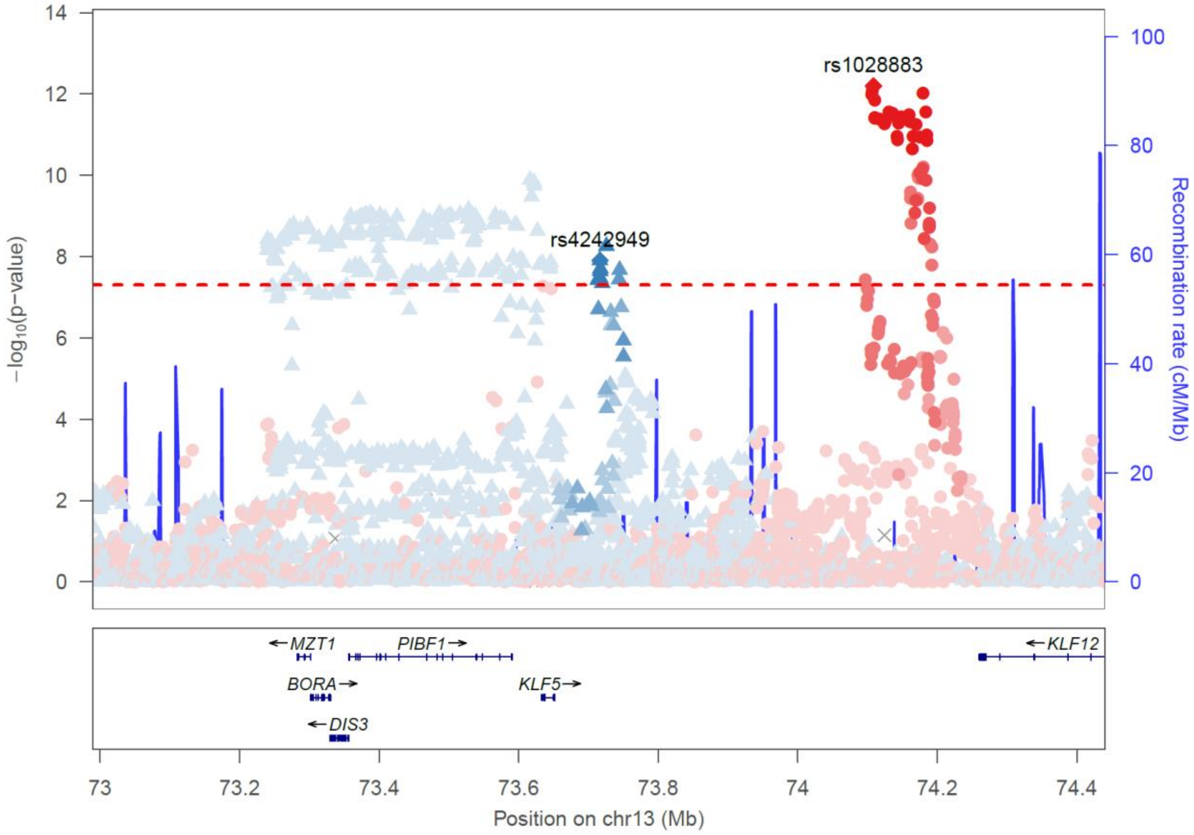

### Locus n27

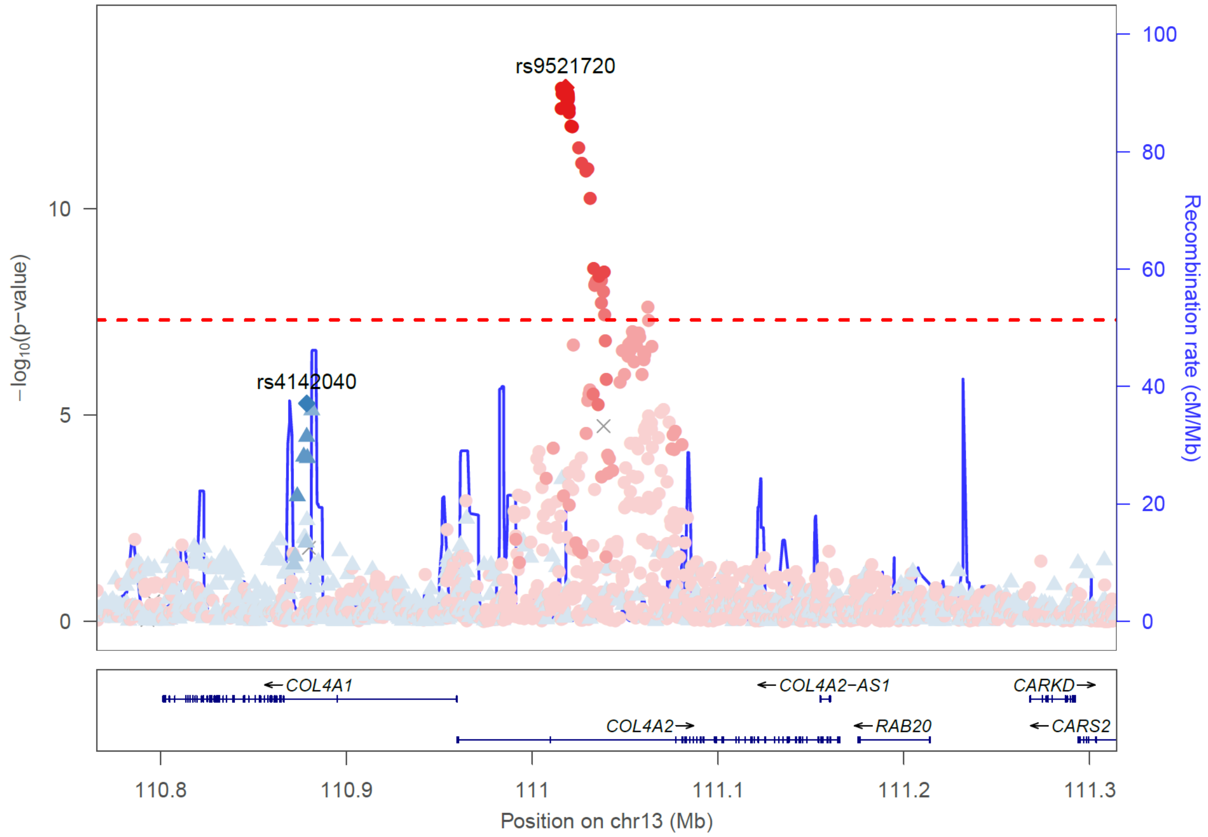


### Locus n29

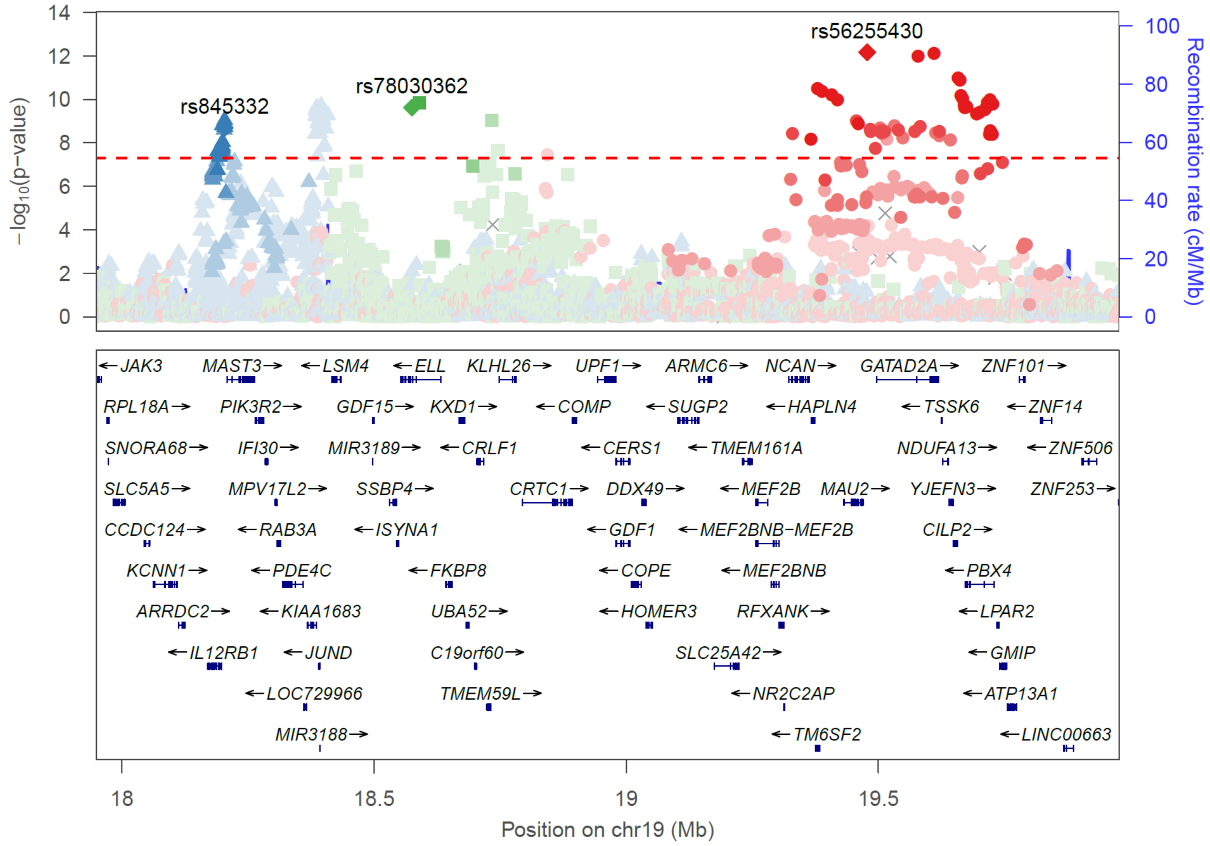

Locus n32

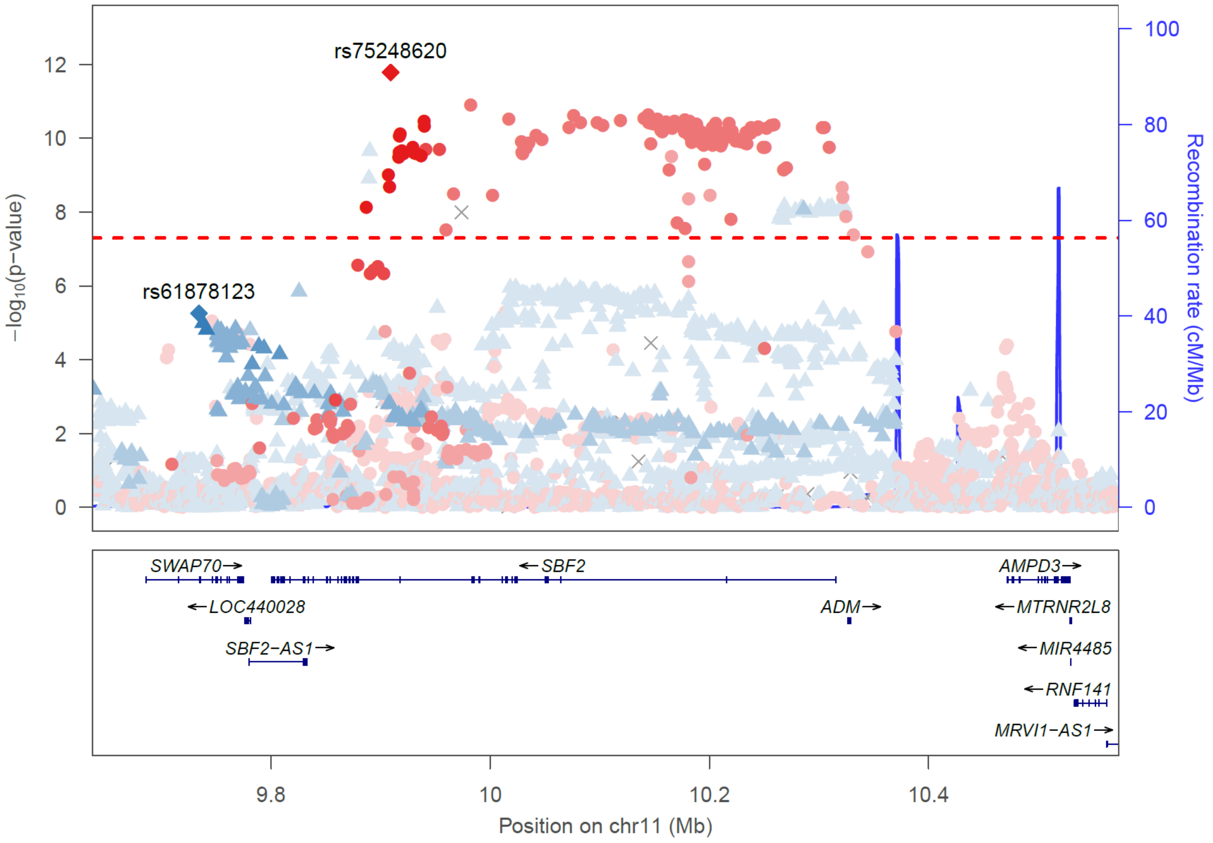


Locus n34

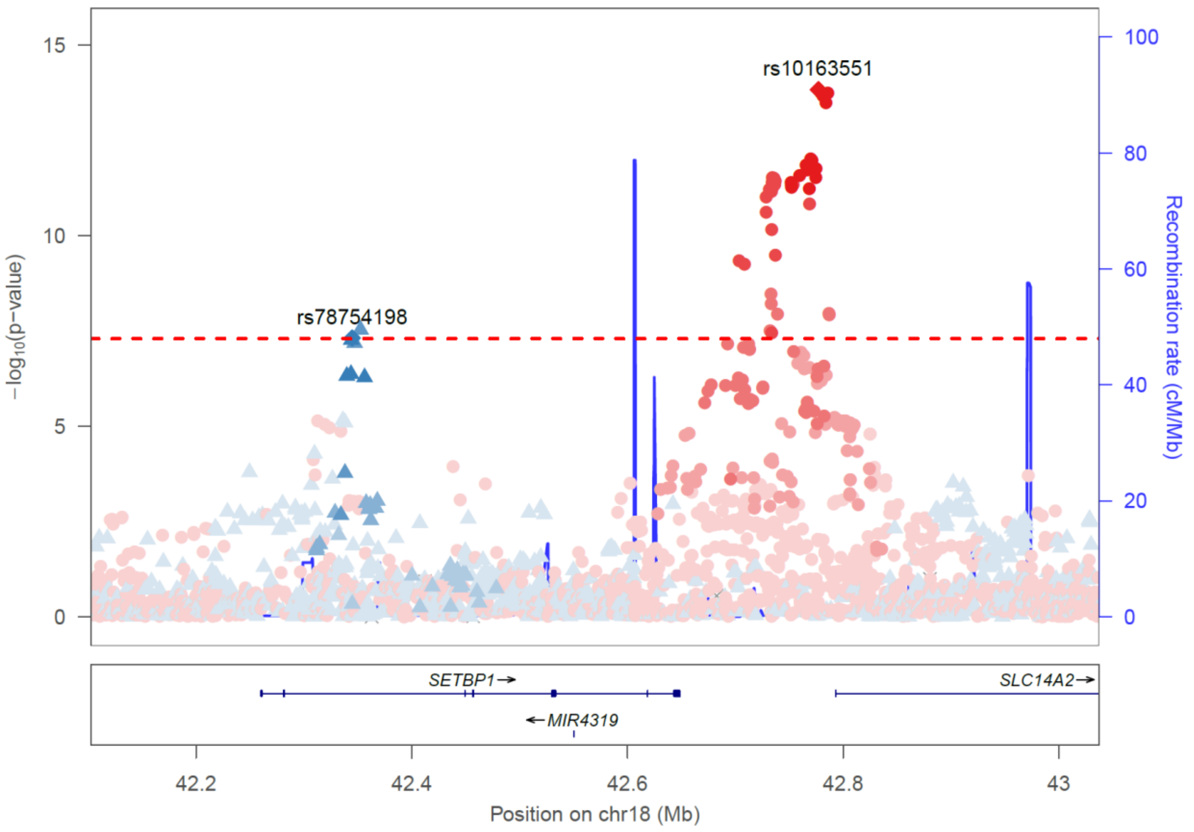

Locus n37

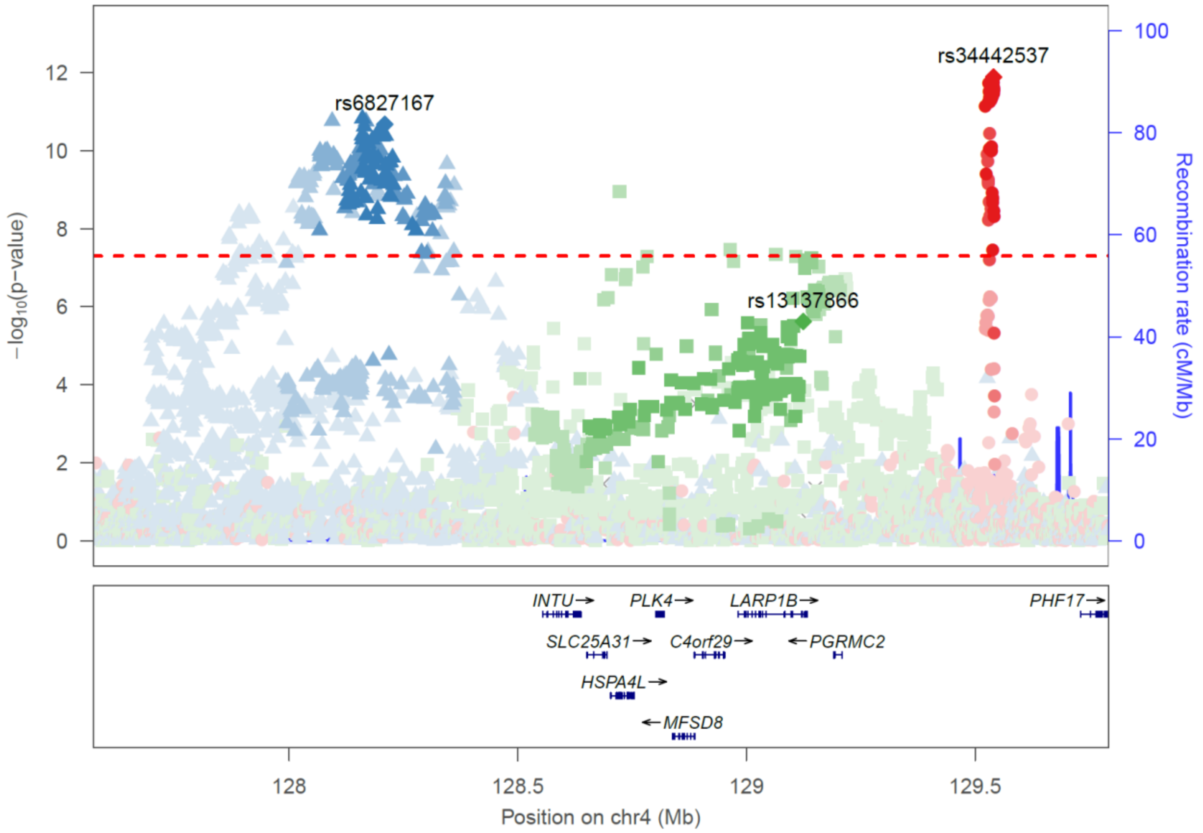


Locus n46

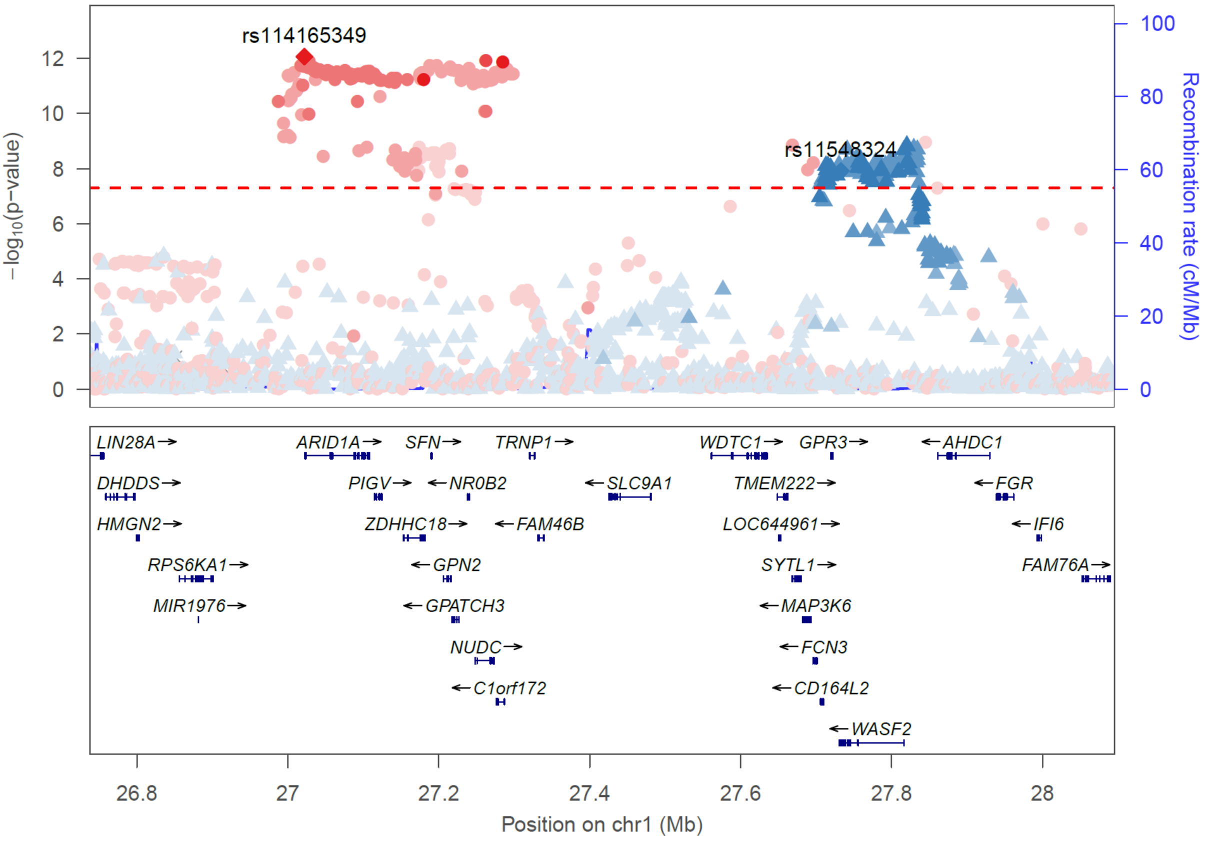

### Locus n47

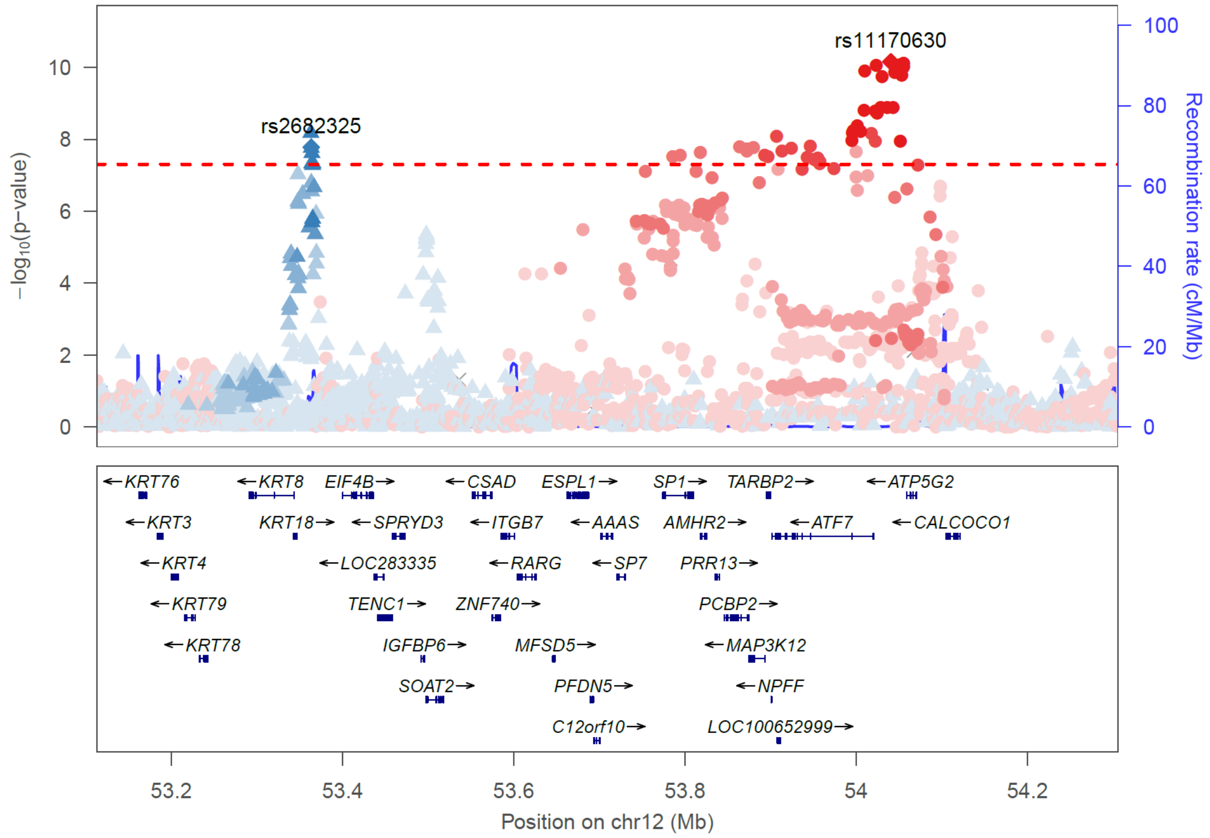


### Locus n55

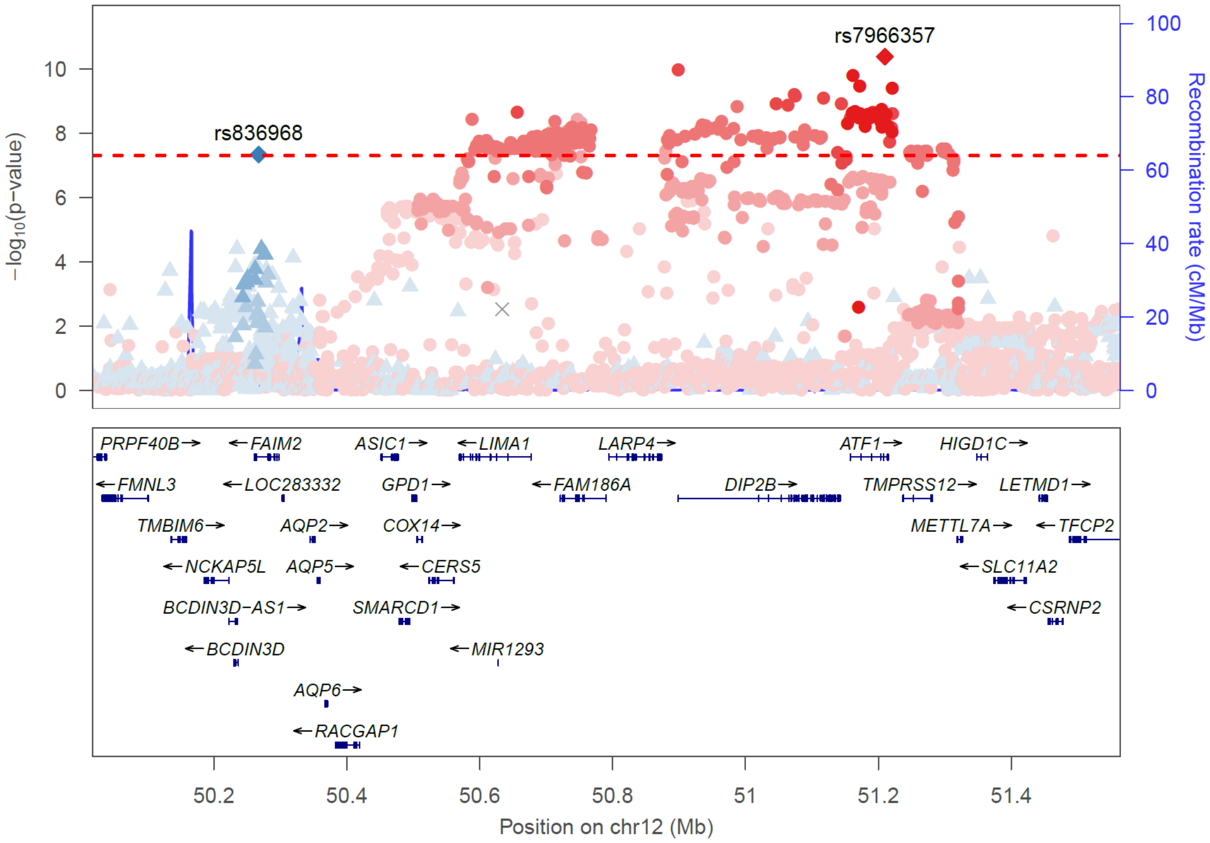

Locus n56

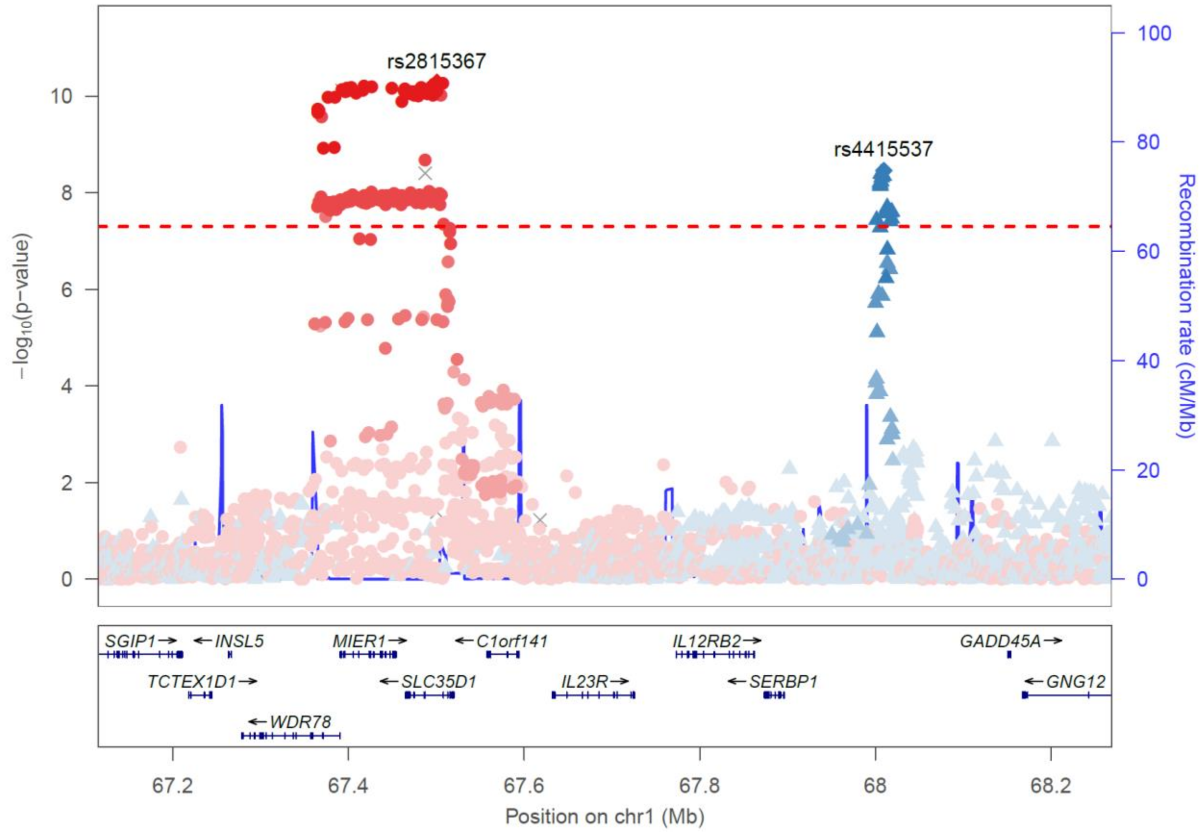


Locus n66

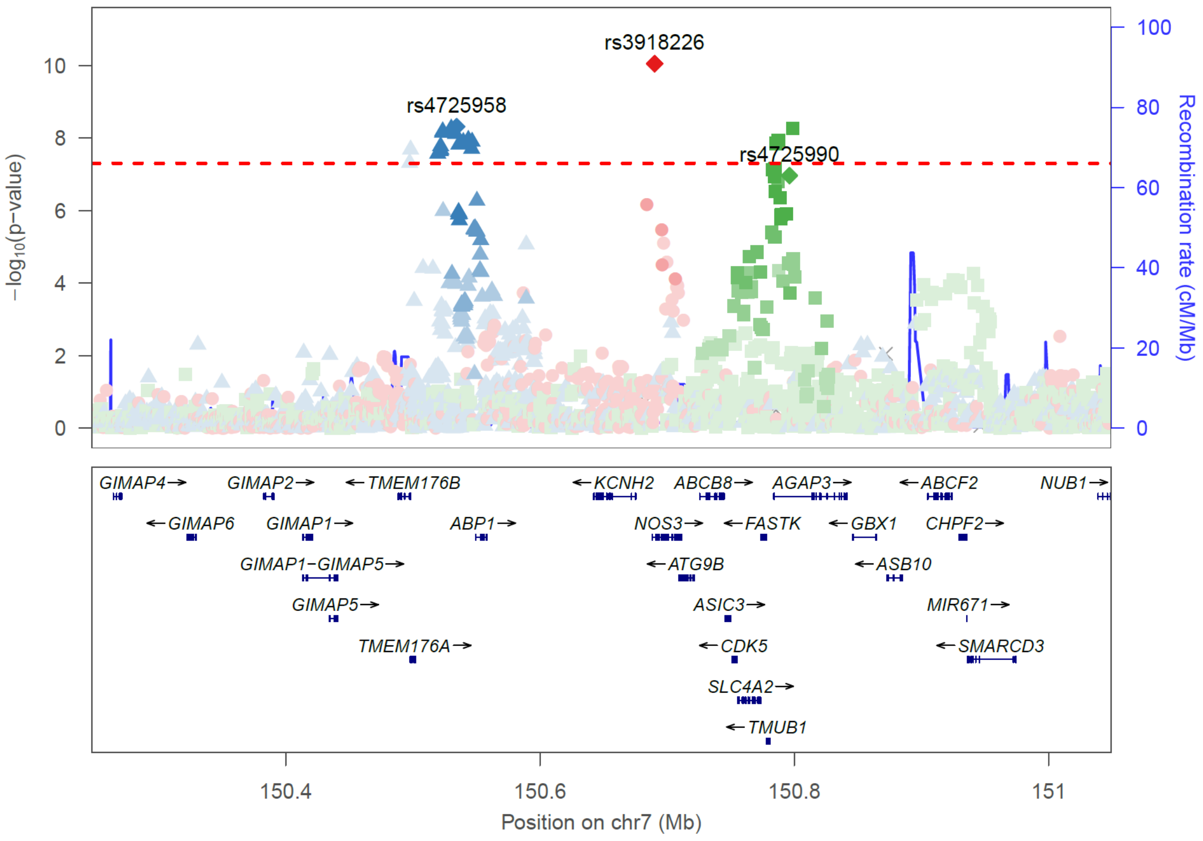

### Locus n76

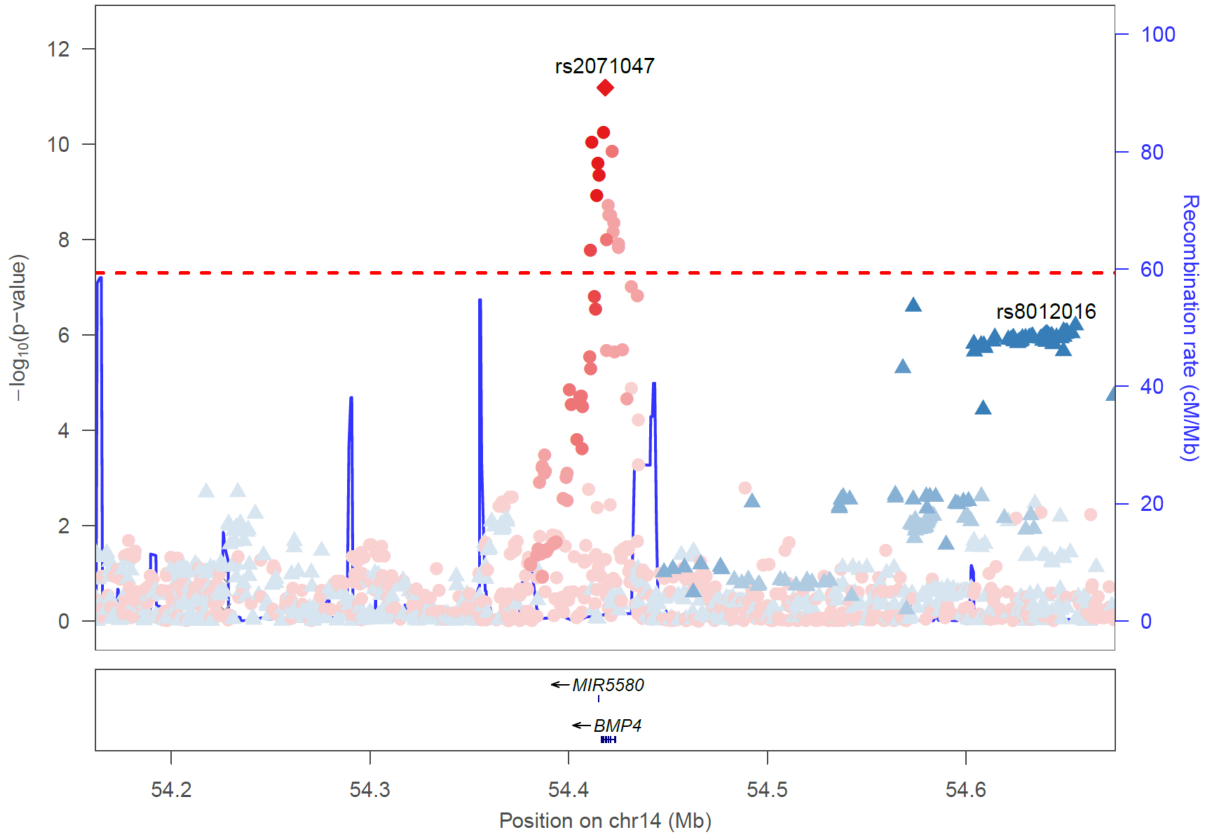


### Locus n80

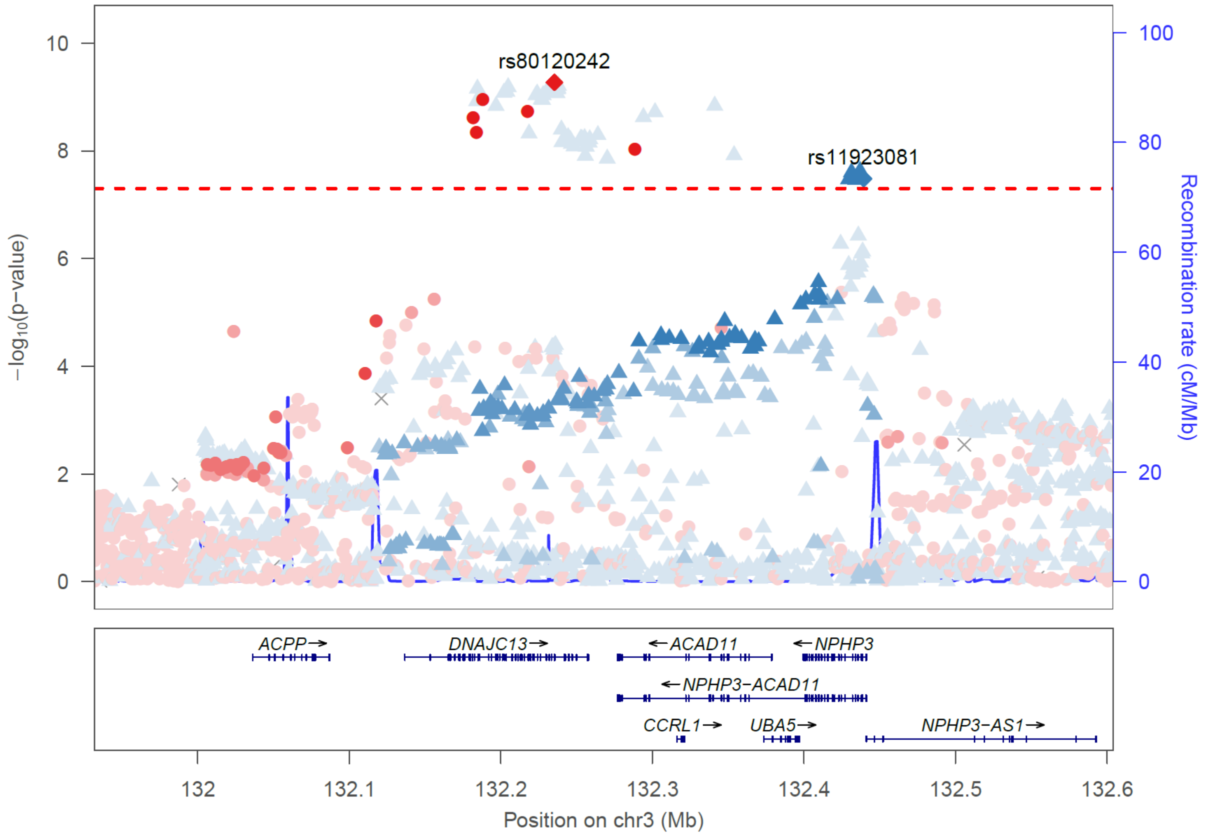

Locus n81

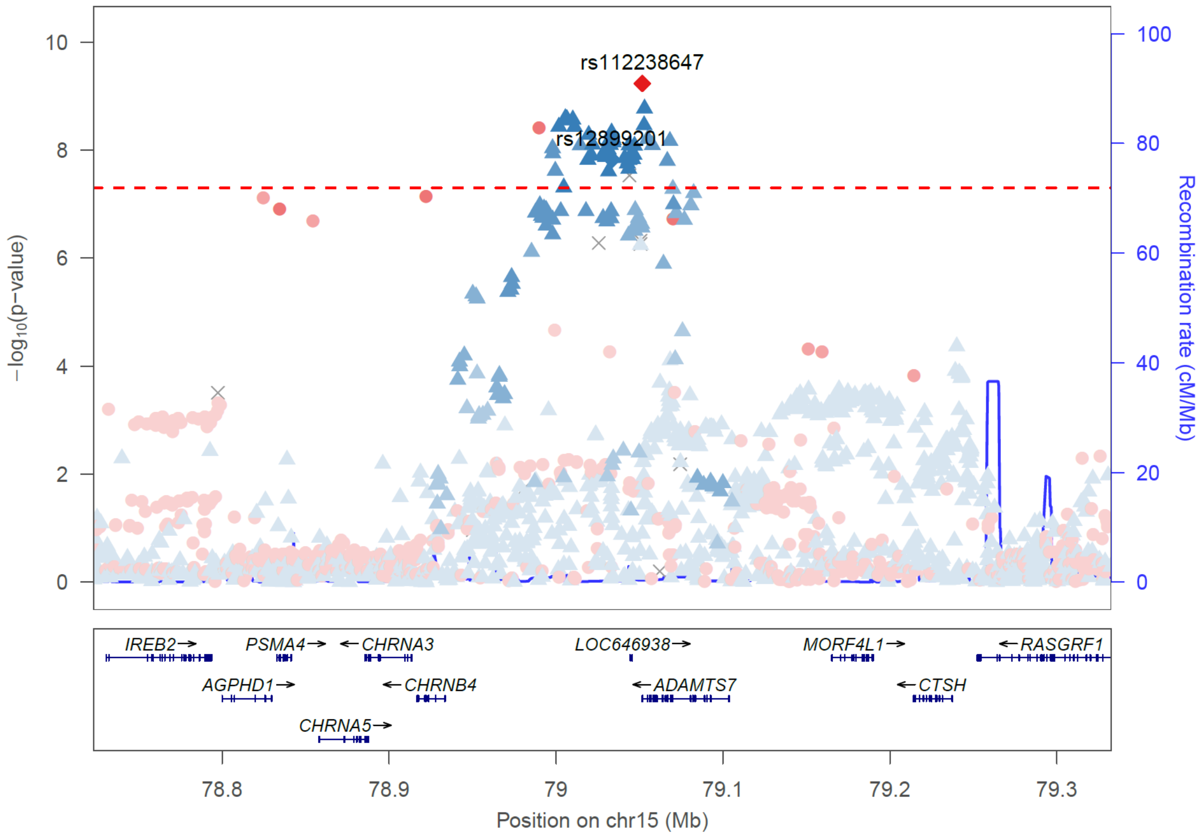


Locus n86

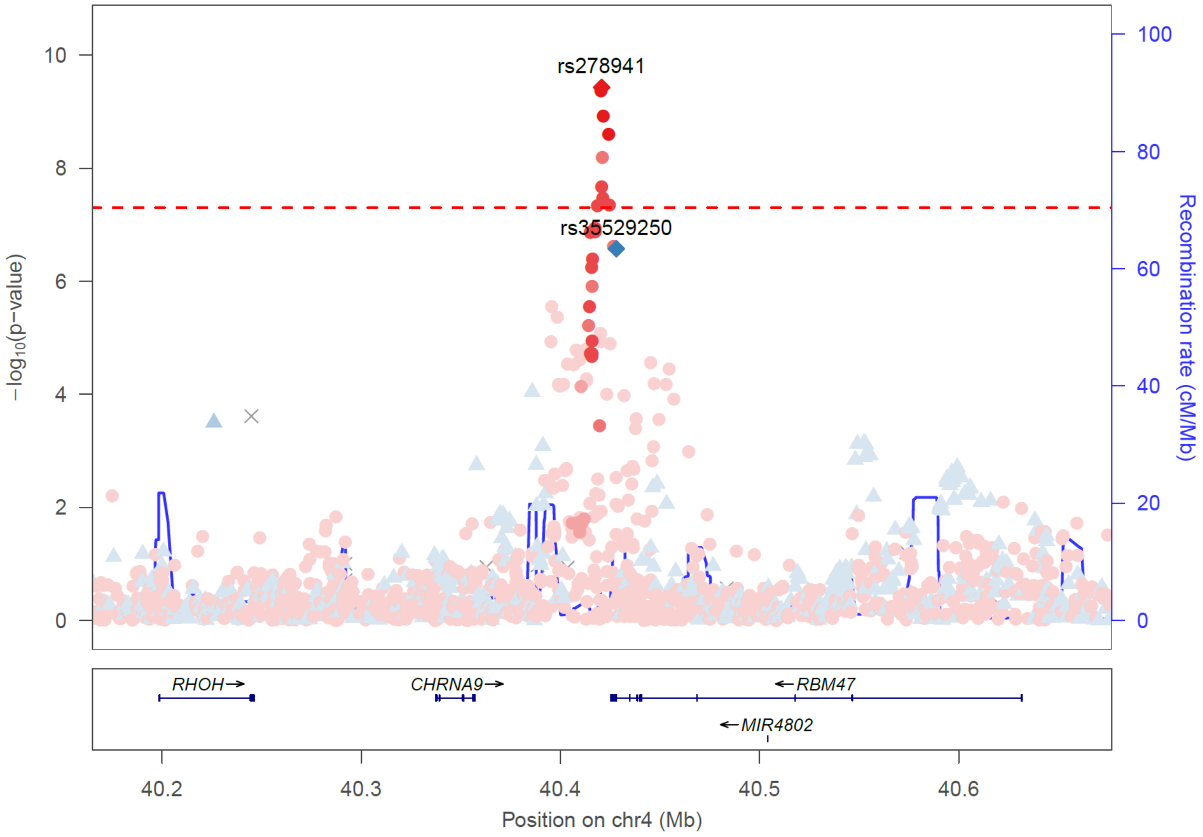

Locus n95

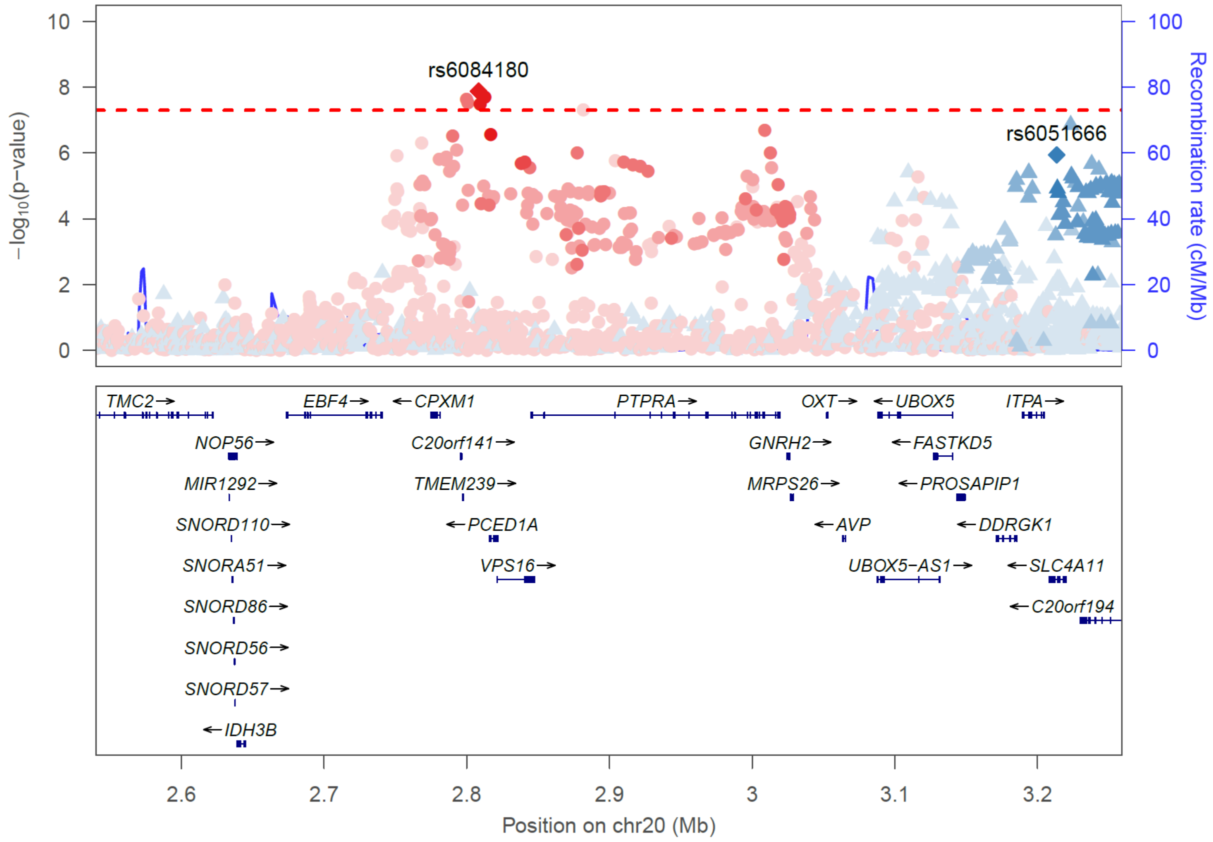


Locus n165

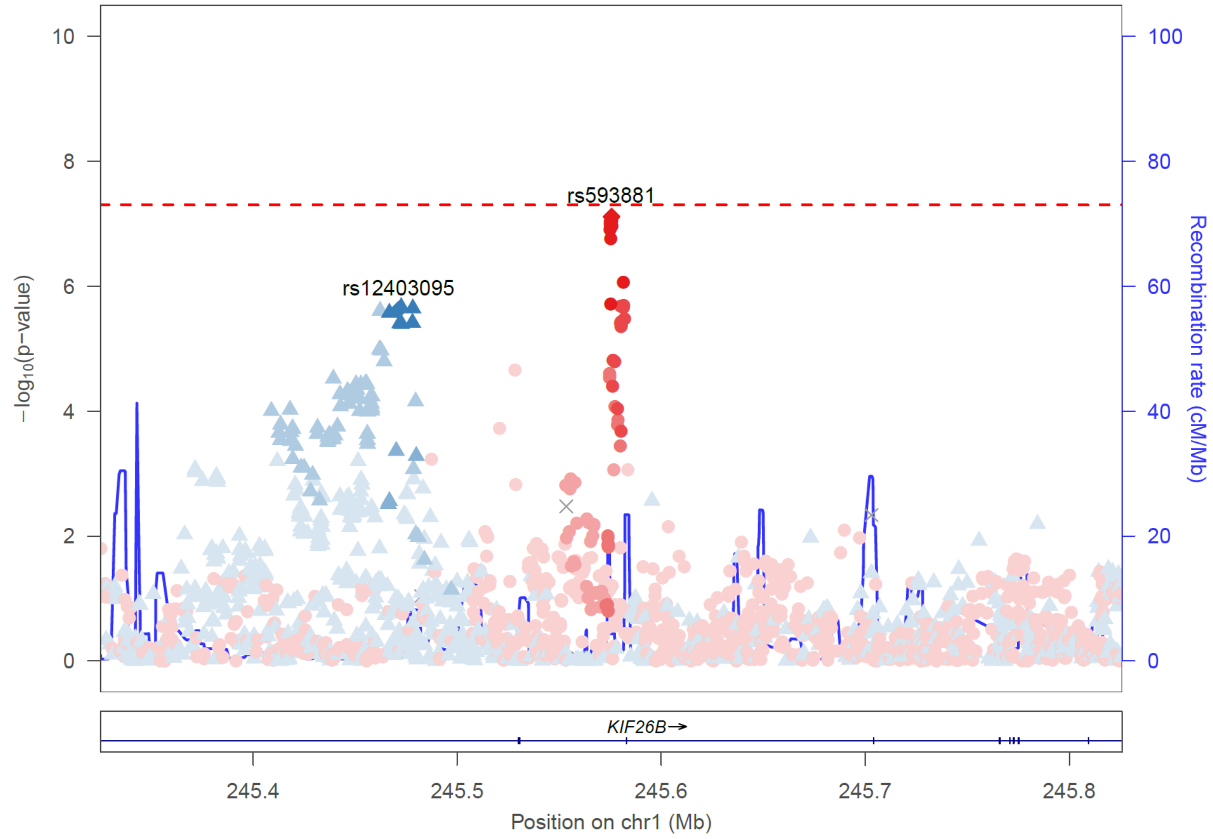

**Supplementary Figure 5. Regional Association Plots (RAPs) at the *UMOD/PDILT* locus.**

The RAPs show association results for eGFRcrea from the approximate conditional analyses with GCTA (n = 1,004,040, European-only meta-analysis). Shown are 4 independent signals observed in the *UMOD/PDILT* locus. For each signal, P-values are conditioned on the other three signal lead variants. Coloring denotes correlation to the respective signal lead variant.

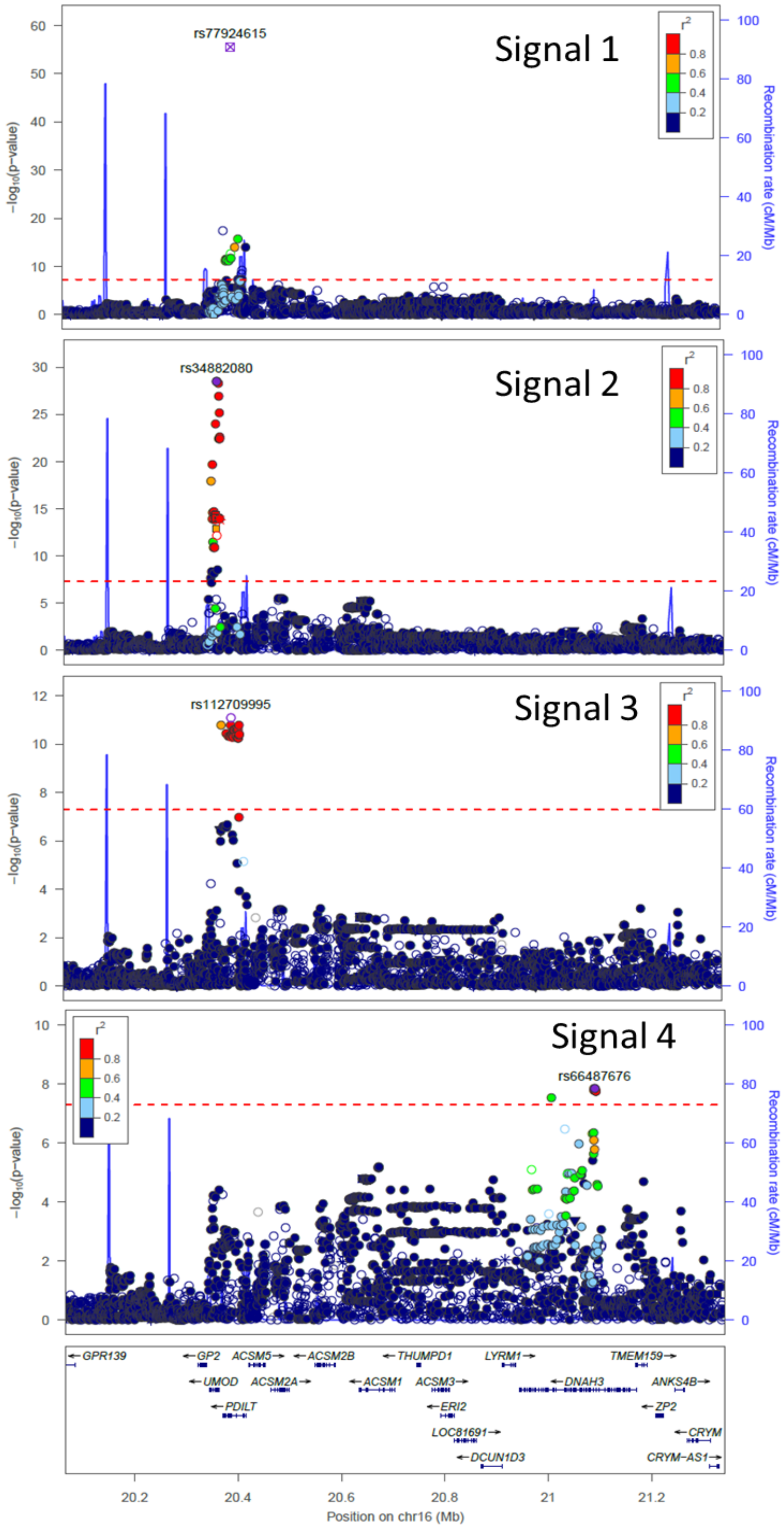

**Supplementary Figure 6. Comparison of Posterior Probability of Association (PPA) between primary lead and fine-mapping variants.**

The scatter plot compares the PPA of the primary analysis lead variants (all-ancestry meta-analysis, n = 1,201,929) with the maximum PPA observed among all credible set variants of the respective signals (based on EUR-only meta-analysis, n = 1,004,040). Orange dots mark all-ancestry lead variants that are not contained in any 99% credible set.

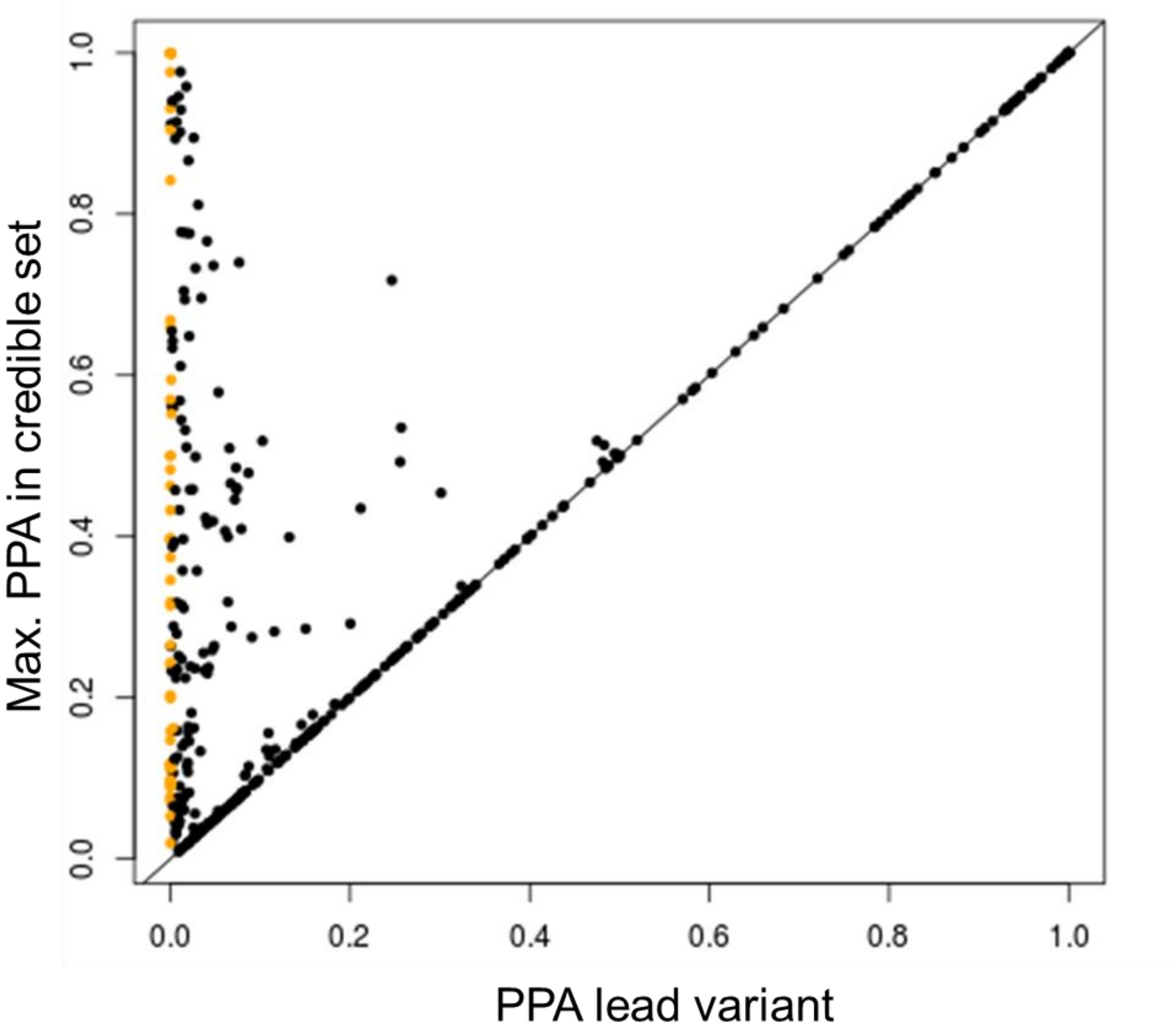

**Supplementary Figure 7. Comparison of colocalization and FDR-based expression analyses.**

The scatter plots compare results from two approaches to evaluate the identified 634 eGFRcrea signals for gene expression effects in two kidney tissues from NEPTUNE [4] (A: tubule-interstitium, B: glomerulus). The first approach is two conduct locus-based colocalization analyses of the effects of variants on eGFRcrea (European-only, n = 1,004,040) and gene expression (two kidney tissues from NEPTUNE). Posterior probability of positive colocalization ($PP_{H4}$) is shown on the x-axis with $PP_{H4}$≥80% denoting 'positive' colocalization between the eGFRcrea signal and the gene expression effects. The other approach is to evaluate the signal's 99% credible variants for significant expression effects. The minimum false-discover-rate (FDR) for gene expression effects observed among the 99% credible variants of the respective signal/gene combination is shown on the y axis, with FDR<5% denoting significant gene expression effects for the variant. Coloring denotes the maximum posterior probability of association (PPA) observed among the respective signals's 99% credible set variants.

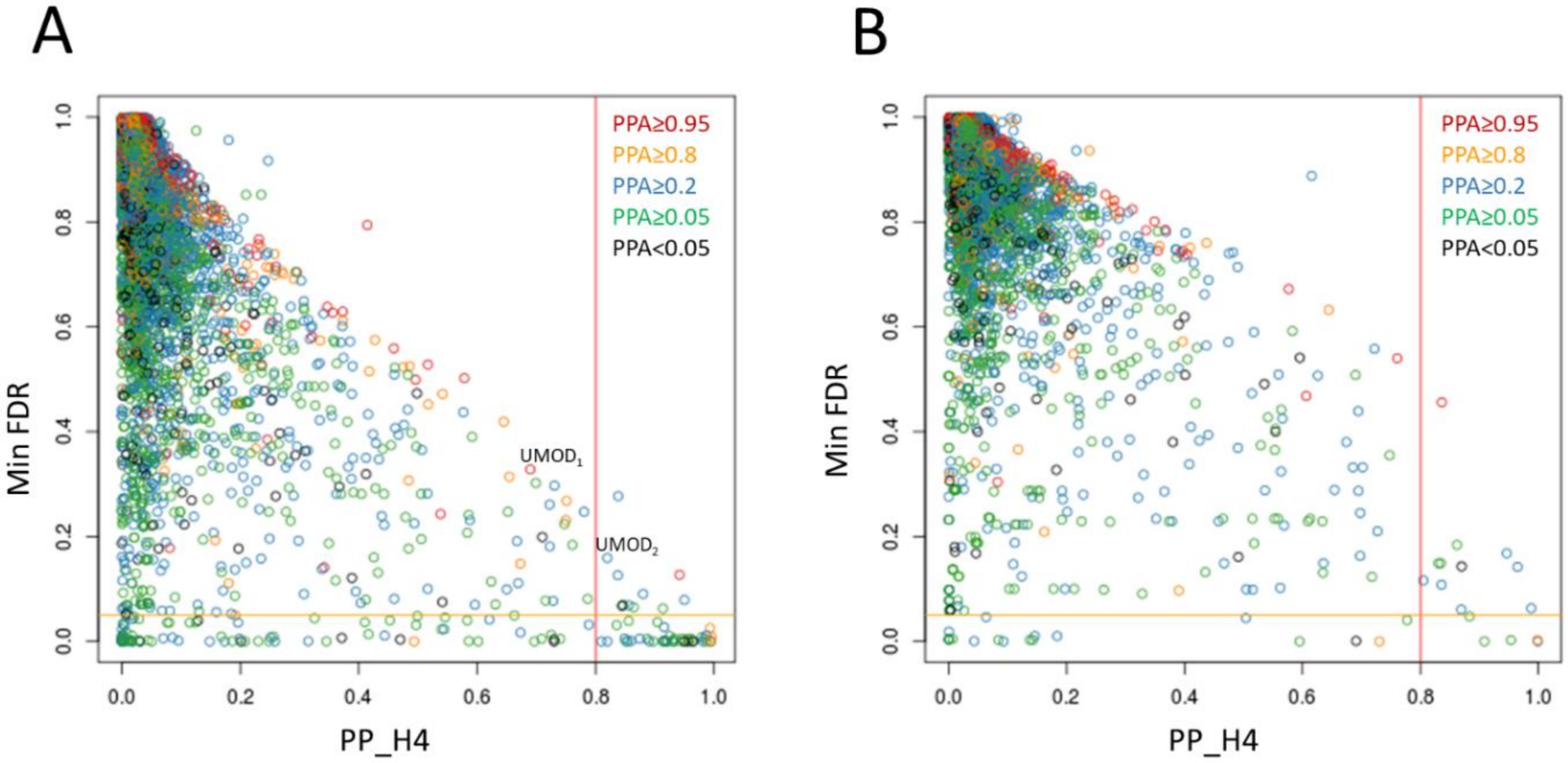

**Supplementary Figure 8. Regional Association Plots (RAPs) for eGFRcrea and ABO expression in NEPTUNE tubule-interstitium.**

The RAPs, obtained with Locuszoom [2], show the associations of genetic variants with eGFRcrea and *ABO* gene expression in tubule-interstitium at the *ABO* locus. Results of the colocalization analysis between eGFRcrea and expression signals are shown in **Table 3**. The RAP helps interpret colocalization results because it allows for a direct comparison between eGFRcrea and *ABO* expression signals. For eGFRcrea, shown are the results from the primary eGFRcrea meta-analysis (n = 1,201,929).

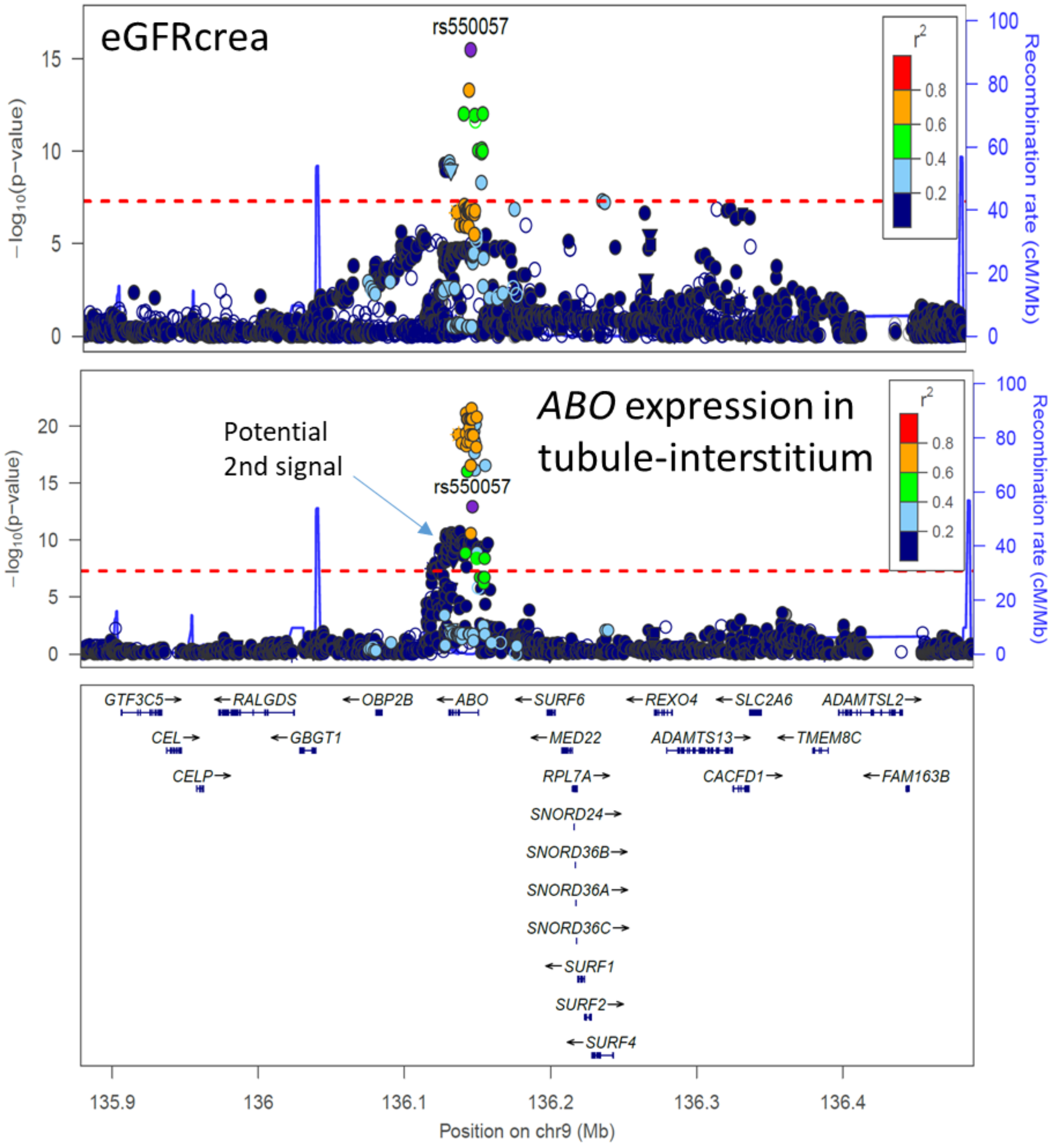
