## Supplementary Note for "Discovery and prioritization of variants and genes for kidney function in >1.2 million individuals"

**Supplementary Note 1. Detailed description of 12 highlighted genes at novel loci.**

From the 2,114 genes underneath the 201 novel loci identified for eGFRcrea (n= 1,201,929), we applied an example weighting in the GPS table (**Supplementary Table 17**). We obtained 12 highly-ranked (gene score >=3) that mapped to novel loci that were validated by eGFRcys or BUN and thus likely relevant for kidney function. The list below shows a brief summary for the 12 genes. Variants with functional consequence are credible variants of the belonging locus with CADD PHRED-like score >15 predicted as non-synonymous:

- *ACOT1:* Encodes for the Acyl-CoA Thioesterase 1 that hydrolyzes acyl-CoAs to the free fatty acid and CoA [1]. No human or kidney relevant mouse phenotypes are described for this gene yet.
- *ACTN4:* Encodes for Alpha Actin 4 that plays a role in tight junction assembly in epithelial cells probably via interaction with MICALL2 [2]. Some rare heterozygous variants are described to cause Glomerulosclerosis, focal segmental, 1 (MIM: 603278).
- *BCS1L:* Ecodes for the mitochondrial chaperon BCS1, which is neccesary for respiratory chain complex III assembly [3]. A spectrum of different missense variants is described to cause Bjornstad syndrome (MIM: 262000), GRACILE syndrome (MIM: 603358), Leigh syndrome (MIM: 256000) or Mitochondrial complex III deficiency, nuclear type 1 (MIM: 124000).
- *BMP4* (rs17563, Val152Ala): Encodes for the secreted ligand Bone Morphogenetic Protein 4, member of the TGF-beta superfamily. It plays an important role in the embryonic development [4]. Heterozygotous mutations in this gene are known to cause microphthalmia with brain and digital anomalies (MIM: 607932) or Orofacial cleft 11 (MIM: 600625). Essential for development of various organs and for neonatal viability in mice [5]. Variants linked to Stickler syndrome and CAKUT in human [6].
- *COL18A1 (rs12483377, Asp1440Asn):* Encodes for the alpha-chain of type XVIII collagen. It seems to be the predominant source of endostatin [7]. Different homo- and heterozygotous mutations are described to cause Knobloch syndrome, type 1 (MIM: 267750) and

Glaucoma, primary closed-angle (MIM: 618880), but kidney related phenotypes were only observed in mouse models.

- *EVC (rs1383180, Arg576Glu; rs2279252, Arg760Glu):* Encodes for the transmembran EvC Ciliary Complex Subunit1, that positively regulates ciliary Hedgehog signaling [8]. Rare homozygous and compound heterozygous mutations are known to cause Ellis-van Creveld syndrome (MIM:225500).
- *HNF1A (rs1800574, Ala98Val):* Encodes for the Hepatocyte nuclear factor 1-alpha, that, as a transcription factor, regulates the expression of multiple genes [9]. Several mutations are described to cause Diabetes mellitus insulin-dependent (MIM:222100, MIM: 612520) and noninsulin-dependent (MIM:125853), MODY typ 3 (MIM: 600496), somatic Hepatic adenoma (MIM:142330) and familiar and somatic Renal cell carcinoma (MIM:144700).
- *INTU:* Encodes for the Inturned Planar Cell Polarity Protein that is part of the CPLANE (ciliogenesis and planar polarity effector) complex which is essential for ciliogenesis [10]. Rare homozygous or compound heterozygous variants can cause Orofaciodigital syndrome XVII (MIM: 617926) and Short-rib thoracic dysplasia 20 with polydactyly (MIM: 617925).
- *LAMC2:* Encodes for the Laminin Subunit Gamma 2, which is involved in attachment, migration and organization of cells into tissue [11]. Rare homozygous variants can cause Epidermolysis Bullosa, Herlitz and non-Herlitz Type (MIM:226700, MIM: 226650), with renal disease as possible comorbidity [12].
- *NPHP3:* Encodes for Nephrocystin 3; part of ciliary proteins required for renal and cardiovascular development [13]. Rare variants can cause Nephronophthisis 3 (MIM: 604387), Meckel syndrome 7 (MIM: 267010) or Renal-hepatic-pancreatic dysplasia 1 (MIM: 208540).
- *NPHS1 (rs3814995, Glu117Lys):* Encodes for Nephrin a member of the immunoglobulin family of cell adhesion molecules that functions in the glomerular filtration barrier in the kidney [14]. A varity of mutations within the coding sequence of this gene is described to

cause Nephrotic syndrome, type 1 (MIM: 256300) which is characterized by proteinuria, hypoalbuminemia, hyperlipidemia, and edema.D.

- *QRSL1 (rs36016898, Ala11Val):* Ecodes a subunit of the Glutaminyl-tRNA Amidotransferase, which catalyses the transamidation of misacylated Glu-tRNA(Gln) in the mitochondria to correctly charged Gln-tRNA(Gln) [15]. Rare homozygous or compound heterozygous variants can cause Combined Oxidative Phosphorylation Deficiency 40 (no kidney phenotype, MIM: 618835).
